# Redefining normal breast cell populations using long noncoding RNAs

**DOI:** 10.1101/2022.09.06.506112

**Authors:** Mainá Bitar, Isela Sarahi Rivera, Isabela Pimentel de Almeida, Wei Shi, Kaltin Ferguson, Jonathan Beesley, Sunil R Lakhani, Stacey L Edwards, Juliet D French

**Author notes:** These two authors contributed equally to this work (Stacey L Edwards, Juliet D French).

## Abstract

Single-cell RNAseq has allowed unprecedented insight into gene expression across different cell populations in normal tissue and disease states. However, almost all studies rely on annotated gene sets to capture gene expression levels and sequencing reads that do not align to known genes are discarded. Here, we discover thousands of long noncoding RNAs (lncRNAs) expressed in human mammary epithelial cells and analyze their expression in individual cells of the normal breast. We show that lncRNA expression alone can discriminate between luminal and basal cell types and define subpopulations of both compartments. Clustering cells based on lncRNA expression identified additional basal subpopulations, compared to clustering based on annotated gene expression, suggesting that lncRNAs can provide an additional layer of information to better distinguish breast cell subpopulations. In contrast, these breast-specific lncRNAs poorly distinguish brain cell populations, highlighting the need to annotate tissue-specific lncRNAs prior to expression analyses. We also identified a panel of 100 breast lncRNAs that could discern breast cancer subtypes better than protein-coding markers. Overall, our results suggest that lncRNAs are an unexplored resource for new biomarker and therapeutic target discovery in the normal breast and breast cancer subtypes.

## INTRODUCTION

Breast cancer is characterized by multiple subtypes, each with distinct molecular features and clinical outcomes. One of the major factors defining these molecular features is their cell-of-origin. Delineating the different cell subpopulations in the normal breast is therefore critical for understanding breast cancer etiology. Breast tumors develop from epithelial cells of the mammary gland, which comprise an inner layer of secretory luminal cells and an outer layer of basal cells with myoepithelial characteristics. Three main epithelial cell populations are known to compose the breast epithelium: basal, luminal progenitor and luminal mature cells (1). While there is evidence for additional subpopulations, including a bipotent progenitor (2), the individual lineages are predominantly self-maintained (1,3). However, the full cellular spectrum and how these contribute to the different subtypes of breast cancer remains to be determined.

Single-cell transcriptomics is allowing us to exploit differential gene expression as a means to define cell types and states. Recent surveys of gene expression in the normal human breast epithelium at single-cell level have discovered new cell types and mapped the trajectory of mammary epithelial lineages (4–7). Like most single-cell RNA sequencing (scRNAseq) analyses, these studies only assess the expression of annotated genes, represented in curated databases such as GENCODE and Ensembl, and sequencing reads that do not map to these regions are lost in the analysis. Since the vast majority of the long noncoding RNAs (lncRNAs) are currently unannotated (8), the expression of lncRNAs in scRNAseq data remains largely unexplored.

LncRNAs are a diverse class of RNA transcripts >200 nucleotides long that lack protein-coding potential. In general, lncRNAs display exquisite tissue- and cell type-specific expression and have been implicated in almost all biological processes. Many lncRNAs regulate nearby gene expression through epigenetic, transcriptional or post-transcriptional mechanisms (9). LncRNAs can also function as molecular scaffolds for protein complexes or promote phase separation of functional subcellular domains (10). LncRNAs also play crucial roles in organ development and establishment of cell lineages by stem-cell fate determination (11). Many lncRNAs are directly implicated in human disease, functioning for example in cancer-related pathways that influence phenotypes such as apoptosis, cell growth, invasion and genomic instability. Here, we discovered >13,600 new lncRNAs expressed in the human breast epithelium and surveyed their expression across epithelial cell subpopulations in very high resolution. We show that lncRNA expression alone can distinguish different cell subpopulations and different cancer subtypes, revealing several novel markers of potential relevance for breast cancer.

## MATERIALS AND METHODS

### Samples, cell sorting and RNA sequencing

Breast tissue samples donated to the Brisbane Breast Bank were obtained from five healthy donors who had breast reduction surgery (12) (Table S1). Reduction mammoplasty samples were processed as described previously (13). In brief, samples were enriched for epithelial cell populations and suspensions were stained with a lineage marker antibody combination (anti-CD31, anti-CD45 and anti-CD140b; Table S1) designed to exclude endothelial, hematopoietic, and leukocyte cells. The remaining lineage-negative cell superpopulation was sorted into basal (EpCAM^low^/CD49f^hi^), luminal progenitor (EpCAM^hi^/CD49f^hi^), and luminal (EpCAM^hi^/CD49f^low^) cell types. Total RNA was extracted and ribodepleted with the Illumina Ribo-Zero Plus rRNA depletion kit. High-quality RNA (RINl’.≥l’.9.0, measured in Agilent 4200 Tapestation) was sent to NovogeneAIT Genomics (Singapore) for 150 paired end short-read total RNA sequencing, performed on the Illumina Novoseq 6000 platform. Each sample was sequenced across 2-3 lanes, yielding 30.6-67.3M read pairs (average of 49.9M). Base call accuracy for every sample was between 99.95-99.99% (average Phred-scale quality between 37 and 39). More details on sequencing quality and general metrics are given in File S1 and Tables S1-4.

### Transcriptome assembly

We designed a multistep computational pipeline for lncRNA discovery from bulk RNAseq based on *de novo* transcript assembly (Figs. 1a and S1 and File S1). The pipeline is subdivided in 3 main steps: (1) pre-processing of reads to remove contaminants (e.g. adapters and reads from ribosomal RNAs) and assess low-frequency k-mers correcting sequencing errors, (2) *de novo* assembly of transcripts and (3) identification of noncoding transcripts and lncRNAs. Additionally, quality control routines were implemented at every step of the pipeline to ensure the overall quality of the assembly met the highest standards (Table S5). All scripts used in the pipeline are deposited in GitHub (Script File 1).

**Fig. 1.**
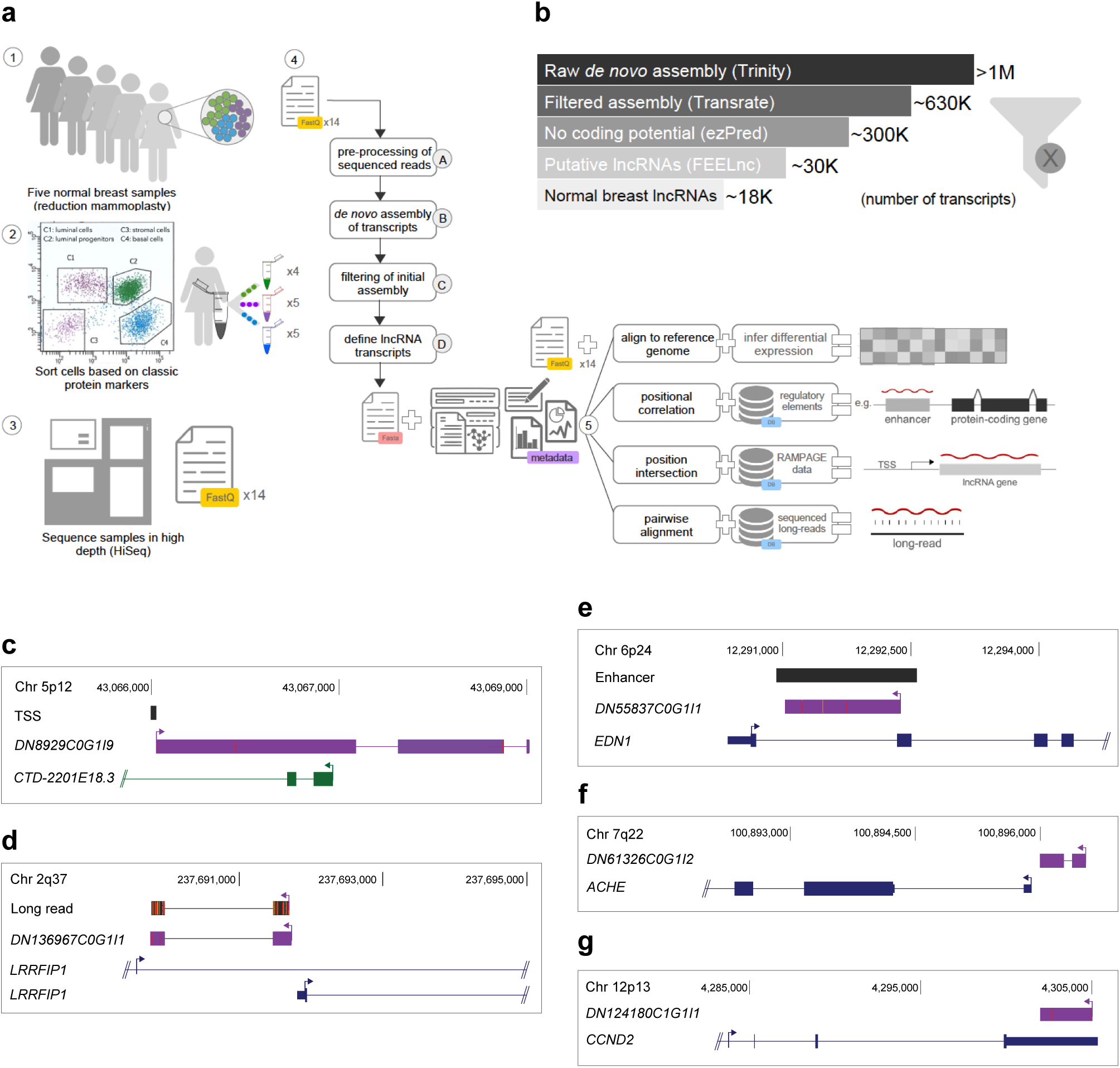
Identification of NB-lncRNAs from human breast epithelium. **a** Schematic of the bulk RNAseq and *de novo* assembly experimental design. Strand-specific RNAseq libraries were prepared from total RNA extracted from FACS sorted breast epithelial cells. A multistep computational pipeline was designed for transcriptome assembly and compared with state-of-the-art tools, showing higher performance. Each main stem (A-D) is described in greater detail in Fig. S1a. **b** Number of assembled transcripts passing each filtering step, from raw transcriptome assembly to identification of NB-lncRNAs. **c-g** UCSC genome browser (hg38) diagram showing NB-lncRNAs (purple) and GENCODE-annotated protein-coding (blue) or noncoding genes (green). Rampage-detected transcription start sites (TSS) and enhancer elements are shown as black boxes.

In step 1, reads were corrected with Rcorrector v.1 (14) and ‘uncorrectable’ reads subsequently removed by FUR-PE v.2016. Adaptors were removed by Trimmomatic v.0.36 (15) and ribosomal RNA reads filtered by BBduk v.2019. Strandness of the RNAseq library was confirmed by RSeQC v.2.6.4 (16). Next, in step 2, we combined reads from all cell populations as input for Trinity v.2.8.4 (17), keeping the normalized read coverage at 50 to prevent fragmented transcripts. Completeness of the initial assembly was confirmed using BUSCO v.20161119 (18). TransRate v.1.0.3 (19) was used to assess assembly quality and as a filtering tool, selecting high-quality transcripts based on features such as read support and quality of base calling. Transcripts were also filtered by explicit read support, using FPKM measures. To differentiate unspliced transcripts from sequencing noise, we enforced a strict read support of 3 FPKM for monoexonic transcripts, relaxing the cut-off to 0.5 FPKM for multiexonic transcripts. We have previously shown that lncRNAs expressed at ∼0.5 FPKM are experimentally confirmed with an 80% success rate (20). In step 3, we identified noncoding transcripts candidates. Four programs (CPAT, CNCI, CPC2 and PLEK, all run through ezLncPred v.1.0 (21)) were used to predict the coding potential of transcripts (Table S6). We considered all transcripts with no coding potential detected by at least two different tools to be true positives. In parallel, we submitted the assembled transcripts to FEELnc v.0.2 (22) for classification. Transcripts were aligned to the GENCODE human genome (version GRCh38) with Minimap2 v.2.16 (23). Both GENCODE-annotated protein-coding and confirmed lncRNA genes were supplied to train the machine learning algorithm of FEELnc_filter_, which filtered non-lncRNA transcripts and transcripts shorter than 200 nucleotides. The FEELnc_codpot_ module was used to compute a coding potential score and assign a confidence level to potential ORFs. The FEELnc_classifier_ module was then used to detect the nearest annotated transcript and further classify the lncRNAs based on genomic position respective to protein-coding genes. Finally, we filtered out lncRNAs identified by FEELnc for which lack of coding potential was not confirmed by ezLncPred, obtaining the set of normal breast lncRNAs (named NB-lncRNAs).

### Long-read and TSS support for NB-lncRNAs

Long-read sequencing and RAMPAGE data were used to assess the transcript structure of NB-lncRNAs (Table S7). Publicly available long-reads from MCF7 and MCF10A cells (24) (PRJEB44348 and PRJNA522784, File S1) obtained by direct RNA sequencing in the Oxford Nanopore Technologies platform were obtained through the SRA portal (https://www.ncbi.nlm.nih.gov/sra). In total, there were 32.7M MCF7 reads of 829 nucleotides and nearly 320,000 MCF10A reads of 971 nucleotides. Additionally, we generated 33.1M long-reads from SUM149 breast cancer cells at an average length of 1,184 nucleotides. Total RNA was sent for long-read cDNA sequencing by PromethION to the Garvan Institute Nanopore Sequencing Facility (Australia). All long-reads were aligned to NB-lncRNAs using standalone BLAT v.35 (25), forcing no gaps in high-scoring blocks and with minimum sequence identity of 80% (-maxGap=0 -minIdentity=80), set to account for the characteristically lower accuracy of long-reads. BLAT was allowed to run in each file for 300 CPU hours. Nearly 3.6M MCF10A transcripts, 1.7 billion MCF7 transcripts and 4 billion SUM149 transcripts aligned to NB-lncRNAs. Alignments were further filtered for a minimum of 70% reciprocal coverage and a maximum of 10% indels at aligned regions, in either sequence. To confirm the TSSs of assembled transcripts, public RAMPAGE data of normal breast (ENCODE entries ENCSR909QWB and ENCSR598TAK, File S1) were obtained from the ENCODE project database (https://www.encodeproject.org/rampage). In-house RAMPAGE libraries were sequenced as 150bp paired-end reads, demultiplexed using the icetea library in R (26). Demultiplexed RAMPAGE libraries and RNAseq libraries were aligned to the reference genome (GENCODE GRCh38) supplemented with the NB-lncRNAs using STAR v.2.7.1a (27). RAMPAGE peaks were called using the *call_peaks* script in GRIT v2.0.4.

### RT-PCR validation of NB-lncRNAs

Total RNA was extracted from FACS sorted luminal progenitor, luminal mature or basal epithelial cells from normal breast using the RNeasy Plus Mini Kit (Qiagen). Two micrograms of total RNA was reverse transcribed using SuperScript IV (Thermo Fisher Scientific) and RT-PCR performed using MyTaq DNA polymerase (Bioline). Fifty cycles of PCR were performed in order to identify low expressed transcripts. Primers are listed in Table S7.

### Annotation of NB-lncRNAs in the human genome

We used Gmap v2020-09-12 (28) to align NB-lncRNAs to the reference genome and create a bed-type file of coordinates. Aligned NB-lncRNAs were intersected by Bedtools v.2.29.0 with the GENCODE GRCh38 transcriptome and matches with minimum overlap of 75% and on the same strand were removed from the final set of NB-lncRNAs, unless the GENCODE transcript was annotated as a lncRNA (Script File 2). The same method was used to intersect NB-lncRNAs with the in-house database of known lncRNAs (File S2 and Table S8). Additionally, we co-localized NB-lncRNAs with known enhancer elements, promoter regions and terminal untranslated ends of annotated genes to provide the first layer of functional annotation (File S3).

### Expression of NB-lncRNAs in the main mammary epithelial cell types

To assess the expression of NB-lncRNAs in the bulk RNAseq samples we used Salmon v.1.3.0 (29) (Script File 3). We then filtered transcripts by expression, requiring at least 1 TPM (transcript per million), to obtain the lists of NB-lncRNAs expressed in each sample. Transcripts present in at least 75% of the replicates of each cell type (i.e. four out of five samples for the luminal mature and basal types and three out of four samples for the luminal progenitor type) were deemed ‘consistently expressed’. From the consistently expressed, those present in only one of the three populations are referred to as ‘uniquely consistently expressed’, or population-specific. We then compared the FEELnc-assigned protein-coding partners of these population-specific NB-lncRNAs with the list of 359 markers retrieved from the literature (Table S8).

### Clustering single cells using different gene sets

Data from a C1 Fluidigm scRNAseq experiment containing 867 cells was obtained from the SRA database (6) (PRJNA450409, File S4). Authors collected samples from reduction mammoplasties performed in age-matched caucasian females at post-pubertal and pre-menopausal stage. No information on their parity and menstrual status was available. Sequencing reads were aligned to the references with Bowtie2 v.2.2.9 (30) and analyzed with RSEM v.1.3.1 (31), using the *single-cell-prior* option, to obtain matrices of transcript counts (Script File 4). The same process was repeated with different references: (i) the entire annotated human transcriptome (GENCODE GRCh38), (ii) the set of NB-lncRNA transcripts, (iii) a subset of ‘(i)’ depleted of protein-coding genes and annotated lncRNAs and (iv) a merge of ‘(i)’ and ‘(ii)’. RSEM count matrices were imported to Seurat v.4.0.4 (32) using Melange v.0.1.0. We then performed quality control, excluding cells with less than 900 detected features when protein-coding genes were considered or 300 cells otherwise (File S4), as well as excluding all genes not detected in at least three cells after filtering. Normalization was performed using the *LogNormalize* function with a scale factor of 10,000. Principal component analysis was performed considering the 2,000 most variable features defined with *FindVariableFeatures* using the variance-stabilizing transformation (vst) method. Before proceeding, we defined the dimensionality and resolution for each experiment after assessing Seurat’s ‘straw’ and ‘elbow’ plots and ClusTree v.0.4.4 (33) clustering trees. Specific parameters selected for each clustering experiment are shown in File S4. Following dimensionality reduction, clustering was performed using *FindNeighbors* and *FindClusters* and visualized with the Uniform Manifold Approximation and Projection (UMAP, File S2) method. Clustering of 10x Genomics data from the same reference (6) was performed in a similar way, except counts were obtained with the CellRanger *count* module. The command-lines used to run Seurat and the accessory tools are contained in Script File 5.

### Assessing the tissue-specificity of NB-lncRNAs

Five random samples each from 27 different human tissues were obtained from public data deposited at the SRA database (PRJNA576920) related with the RNA Atlas publication (34). Raw files were processed through our in-house RNAseq analysis pipeline, described at Script File 6 and deposited at GitHub. Briefly, raw sequencing reads were assessed for quality control, trimmed and aligned to the GENCODE human genome (version GRCh38). Transcript abundances were estimated in expected counts and summarized as TPM values. Transcripts were classified as either protein-coding, annotated lncRNAs or NB-lncRNAs. Expected counts were transformed to log-scale and quantile-normalized using in-house scripts. Tispec (https://github.com/roonysgalbi/tispec) was used to calculate tissue-specificity per transcript. For every transcript, the tissue with highest tau value was recorded and the occurrences of each tissue were summarized as the global tissue-specificity. The same process was repeated for all three subsets of transcripts (protein-coding, annotated lncRNAs and NB-lncRNAs).

### Clustering brain cells based on NB-lncRNA expression

The transcriptomes of 466 brain cells were obtained from the SRA database (35) (PRJNA281204, File S2). Cells were clustered using the protocol described above, with at least 300 genes required per cell and a minimum of three cells expressing each gene. For comparison, we also clustered cells using the expression of GENCODE-annotated genes, raising the minimum number of required genes per cell to 900. We used labels provided by the authors and the marker genes listed in the original publication (35) to characterize the cell subpopulations in each cluster.

### Cluster specificity index

An in-house method (Script File 6) was used to compute gene expression averages across cells in each cluster (with R v.3.6.2). Briefly, specificity indices for each transcript were computed using TSPEX v.0.6.1, a package offering twelve distinct tissue-specificity metrics (36). After consideration, we selected the TSI (37) as our preferred metric of specificity. The TSI defines specificity as the median expression of each transcript in each cluster divided by the sum of the medians in all clusters: 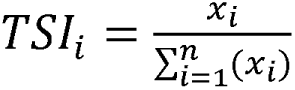

Cluster specificity indices (CSI) were then computed by averaging the specificity indices of all transcripts per cluster and multiplying the resulting number by a factor of 100, for easier interpretation.

### Cluster stemness

Teschendorff and Enver developed a method named Single-Cell ENTropy (SCENT) to quantify the potency of single cells (38). We used SCENT to calculate the signaling entropy (SR) of cells which is a measure of gene expression promiscuity based on a gene interaction network. To account for the contributions of lncRNAs, we replaced the in-built protein-coding gene interaction network of SCENT with an in-house coding <-> noncoding network based on LncRNAs2Pathways (39) data. Details on how we obtained information from LncRNAs2Pathways, formatted the network and incorporated into SCENT are available in File S5. Cell-level SR scores were calculated based on gene expression matrices and averaged to yield cluster-level SR scores for L-clusters (Script File 7). Cluster-level SR values were interpreted as a measure of stemness (i.e. the higher the value, the more likely for a cluster to harbor stem-like cell subpopulations). Additionally, we used Monocle v.2.18.0 (40) to analyze the distribution signaling entropy along reconstructed cell hierarchies (Script File 8). Monocle applies a negative binomial model to test for differential expression using the same gene expression matrix (of raw counts) provided to Seurat. After normalizing the data and filtering for extreme RNA numbers, we used Monocle’s *reduceDimensions* and *orderCells* functions to perform dimensionality reduction with the DDRTree method and sort cells along the inferred trajectory, coloring by cell-level SR value.

### Differentiation trajectories in the mammary epithelium

Being confident that the obtained cell hierarchies recapitulated the mammary differentiation process, we used Slingshot v.1.8.0 (41) to plot trajectories, based on Seurat clustering and manually assigning the root state to cluster L3 (Script File 9).

### NB-lncRNA signatures to discern breast cancer subtypes

We used an in-house pipeline (Script File 10) to map RNAseq reads from TCGA (42) to the GENCODE human transcriptome supplemented with NB-lncRNAs. Briefly, we used Trimmomatic v.6.36 to remove sequence contaminants, STAR v.2.7.1a (27) for sequence alignment and RSEM v.1.3.1 (31) to estimate read counts per transcript, adding TPM values to compute gene expression levels. Subtypes were assigned to each TCGA sample according to the classification in Netanely *et al* (43). Bulk expression was defined as the average across all samples of each subtype for TCGA data and average pseudobulk expression for normal breast cell clusters. Correlation tests between bulk expression profiles of A-clusters or L-clusters and TCGA subtypes were based on Pearson coefficients and associated p-values, calculated in R. The Genefu package v.2.22.1 (44) was used to compare bulk expression profiles of annotated genes in clusters with molecular subtypes of breast cancer defined by either the ‘PAM50’ or the ‘ClaudinLow’ panels.

To find markers of each breast cancer subtype, MGFR (Marker Gene Finder in RNAseq data) v.1.16.0 (45) was run independently 500 times with sets of twelve TCGA samples (three samples per subtype) in a bootstrap manner (Script File 11). Each run resulted in a list of candidate markers and the 300 most frequently retrieved genes were selected for the final set of subtype markers. We repeated the same process for GENCODE-annotated genes and NB-lncRNAs separately. To assess the performance of the method, we compared the marker lists obtained for annotated genes with previously published marker lists (46), calculating the significance of the overlap with Fisher’s exact tests (using the *GeneOverlap* R function, Script File 12). The top 20 most frequent (from the 300) NB-lncRNA markers were used to generate PCA plots and assess their ability to discern between different TCGA subtypes.

### Statistical analyses and data plotting

When two sets of annotated genes were being compared for enrichment, we used the *GeneOverlap* function of R, with background defined as the total number of annotated genes (unless otherwise stated) and used Fisher’s exact test to define significance. R was also used to assess normality of datasets, perform T-tests, Wilcoxon tests and calculate Pearson correlations and compute their associated p-values. To calculate the average, median, sum, minimum and maximum values of any set of values, we used an in-house script. We used R functions *ggplot*, *heatmap* and *prcomp* to plot most graphs, including boxplots, bar plots, pie charts, heatmaps and PCA plots. Unless otherwise specified, R v.4.0.2 was used. The main accessory scripts are provided in Script File 12.

## RESULTS

### Assembly of the normal human breast transcriptome

We obtained normal breast tissue from five healthy women undergoing reduction mammoplasty (Table S1), selected epithelial cells based on the absence of CD31, CD45 and CD140b markers (Table S1) and sorted these cells into basal, luminal progenitor and luminal mature populations, according to their levels of EpCAM and CD49f (Table S1). Bulk strand-specific RNA sequencing was performed on each sorted population at high depth, to allow accurate transcript assembly and assignment, even for genes with very low expression levels (17). On average ∼50M reads were sequenced per cell population per sample (Table S2). Base call accuracy surpassed 99.99% after trimming and correction (Tables S3 and S4). *De novo* assembly of transcripts and discovery of lncRNAs were conducted using an in-house pipeline (Fig. 1a, Fig. S1a and File S1), which, according to the metrics defined in a recent benchmarking study (47), performed >10% better than the best-ranked assembly tool (Table S5; File S1).

The initial assembly contained >1M preliminary transcripts (Fig. 1b) and covered one-third of the human genome (Table S5). Completeness was assessed with BUSCO (18), confirming the presence of >99% of the conserved eukaryotic orthologs (Table S5). We optimized the initial assembly with TransRate (19), filtering transcripts with lower scores. The optimized assembly contained 627,743 high-quality transcripts (average contig length of 826 nucleotides and N50 of 1,383; Fig. 1b and Table S5). To ensure the authenticity of the assembled monoexonic transcripts, we set a strict expression level cut-off (3 fragments per kilobase per million, FPKM), six times the minimum required for multiexonic transcripts (File S1). Approximately 95% of multiexonic (∼85,000) and 60% of monoexonic (∼300,000) transcripts were retained after enforcing minimum expression support. The complete transcriptome of the normal breast epithelium consisted of 384,182 coding and noncoding transcripts. General gene expression profiles, length, exon numbers and splice site motifs of NB-lncRNAs were in agreement with previously annotated lncRNAs, supporting their status (File S1).

### Discovery and annotation of lncRNAs in human breast epithelium

We used ezLncPred (21) to run multiple predictors (CNCI, CPAT, CPC2 and PLEK) and detected coding potential in ∼20% (83,034) of the transcripts (Fig. 1b, Table S6). In parallel, we used FEELnc (22) to classify transcripts as lncRNAs, a machine learning-based algorithm which performs equal to or better than GENCODE and NONCODE consortia classifiers (48,49). FEELnc identified 30,722 lncRNA candidates (Fig. 1b), 90% of which were supported by two or more ezLncPred predictors (Table S6). To minimize false positives, we filtered out the 10% unsupported lncRNA candidates, then discarded candidates with >75% overlap to annotated protein-coding transcripts on the same strand. The final set of curated lncRNAs contained 18,364 transcripts (Fig. 1b), which we will refer to as normal breast lncRNAs (NB-lncRNAs).

We tested the performance of our transcript reconstruction strategy by assessing the full-length read support and transcription start site (TSS) of the assembled NB-lncRNAs. Using public RAMPAGE data of normal breast samples (50) and in-house RAMPAGE data of breast cell lines (BT549, MCF10A, MDAMB231 and SUM149), we confirmed the TSS of 10% (1,810) of the NB-lncRNAs (Fig. 1c). Additionally, full-length transcripts from public long-read sequencing of MCF7 and MCF10A and in-house long-read sequencing of SUM149 breast cells confirmed the exon structure of 7% (1,310) of the NB-lncRNAs (Fig. 1d). Overall, we confirmed critical features of >15% (2,863) of NB-lncRNAs (Table S7 and File S1), in agreement with previously reported observations (e.g. (51,52)). We also used RT-PCR to validate NB-lncRNAs expressed in luminal progenitor, luminal mature or basal epithelial cell populations (defined by cell sorting). Twenty-five of the thirty NB-lncRNAs were validated at the expected size (Fig. S1b).

From the 18,364 NB-lncRNAs, 3,642 coincide (i.e. at least a 75% bidirectional match) with GENCODE-annotated transcripts and 1,116 with ncRNAs from our in-house compilation of public databases (Table S8; File S2). These add to >20% of all NB-lncRNAs. Moreover, partial overlap (i.e. at least a 50% unidirectional match) to annotated noncoding isoforms was detected for another 20% of the NB-lncRNAs, which are likely novel isoforms of known genes (File S1). GENCODE-annotated NB-lncRNAs include 535 antisense genes, 388 intergenic lncRNAs and 100 pseudogenes (Table S9). In total, >85% of the NB-lncRNAs are novel or known lncRNAs and the remaining are noncoding isoforms of annotated protein-coding genes (Table S9). Notably, transcripts of >100 breast cancer-related genes from lncRNAfunc (53), Lnc2Cancer (54) and a study by Diermeier et al (55) were recovered as NB-lncRNAs (Table S9; File S3).

Many lncRNAs act by regulating the expression of nearby protein-coding genes. We characterized regulatory NB-lncRNAs by reciprocal sequence overlap, identifying transcripts originating from known enhancer elements (enhancer-derived lncRNAs, elncRNAs), promoter regions (promoter-associated noncoding RNAs, pancRNAs) or terminal untranslated ends (terminus-associated lncRNAs, TALRs) of annotated genes. In total, we detected 349 NB-elncRNAs, 1,968 NB-pancRNAs and 825 NB-TALRs. Details are provided in the Supplementary Material (File S3; Table S10) and examples of each class are shown in Figs. 1e-g.

### Bulk expression of NB-lncRNAs in normal breast epithelium

We investigated the expression patterns of the NB-lncRNAs in the sorted breast cell populations used for the *de novo* assembly. On average, 10,000-12,000 transcripts are expressed at >1 TPM in each sample, with 65-75% being expressed in samples of the same cell population from all five individuals (Table S11). Over one-third of the expressed transcripts are common to the three cell populations and, as expected, the two luminal cell populations share nearly four times more expressed transcripts with each other than with the basal population. Considering only transcripts expressed at >1 TPM in at least 75% of the individuals, we found 6,371 NB-lncRNAs in luminal mature cells, 5,922 in luminal progenitor cells and 4,577 in basal cells. From these, respectively 1,424 (22%), 957 (16%) and 752 (16%) are unique to each cell population and these population-specific transcript sets are enriched (p-values of overlap <= 7.2e-62) in NB-lncRNAs located nearby annotated markers of the respective cell population (56) (Figs 2a-c). These results provide evidence that expression of population-specific NB-lncRNAs may be related to annotated marker gene activity in the corresponding cell types.

**Fig. 2.**
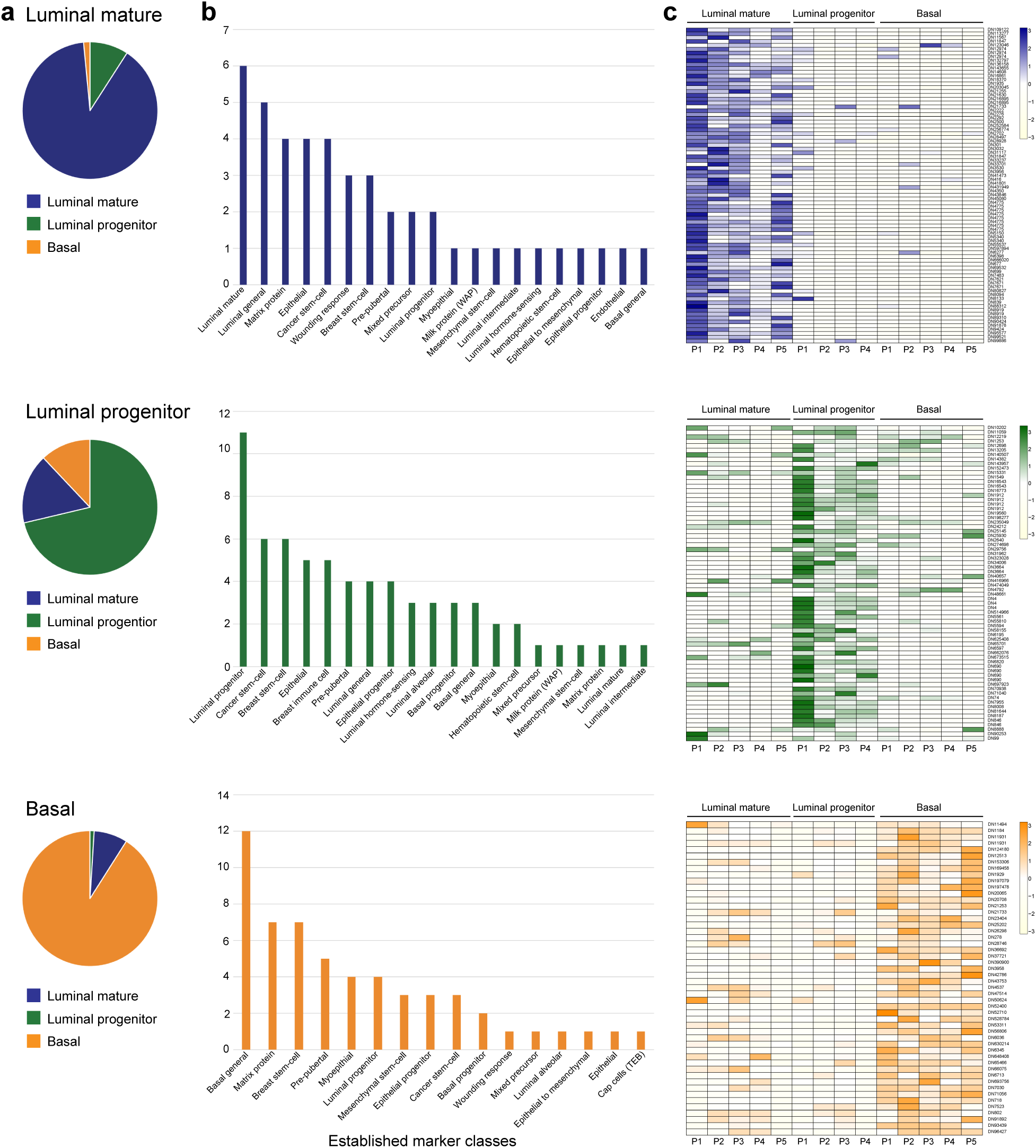
Cell population-specific NB-lncRNAs target protein-coding markers of the same cell type. **a** FEELnc-assigned partners of NB-lncRNAs specifically expressed in each of the three main epithelial cell populations [i.e. luminal mature (top, blue), luminal progenitor (middle, green) and basal (bottom, orange)] were compared with previously reported markers of cell types, showing significant overrepresentation of the corresponding type. Enrichment was confirmed based on the p-value obtained for Fisher’s exact tests. **b** Similar analyses were performed using an in-house dataset of known markers of several normal breast cell types, also showing a higher proportion of the population-specific NB-lncRNA partners are characteristic of the corresponding cell type. **c** Heatmaps of gene expression confirm the NB-lncRNAs as population-specific.

FEELnc identifies the closest annotated genes to the novel lncRNAs and predicts the most likely mRNA-lncRNA partners based on the lncRNA class (22). The term ‘partners’ refers to lncRNA-mRNA pairs predicted by FEELnc. Among the FEELnc-assigned partners of the population-specific NB-lncRNAs for luminal mature, luminal progenitor and basal cells, respectively 27, 34 and 31 are established markers (File S2) of normal breast cell populations (Tables S8 and S11). FEELnc-assigned partners of population-specific NB-lncRNAs in the basal cell population include known basal markers, such as *ACTA2*, *CCND2*, *DKK3*, *ITGA6*, *SPARC*, *TP63* and *VIM* (Table S11). Similarly, in the luminal mature and luminal progenitor cell populations, partners of several population-specific NB-lncRNAs are established marker genes, including luminal mature markers *AKT1*, *ANKRD30A*, *IGF1R*, *PRLR* and *SYTL2* and luminal progenitor markers *CD24*, *CD44*, *KIT*, *LTF*, *PROM1*, *SAA2* and *SLPI* (Table S11). Notably, several NB-lncRNAs are expressed in both luminal populations (but not in basal cells), for example DN34862C1G1I1, an antisense transcript of ELF3 which is a general marker of luminal cells (57).

### NB-lncRNA expression distinguishes the main breast epithelial cell types in scRNAseq data

To explore the ability of NB-lncRNAs to distinguish normal breast cell subpopulations, we quantified the expression of NB-lncRNAs using published scRNAseq data of 867 cells from three donors, with >1.5M reads per cell (6). Using flow cytometry, Nguyen *et al* sorted cells from the luminal and basal populations before performing scRNAseq (File S4). On average, each cell expressed close to 900 NB-lncRNAs, with over one-third expressing at least 1,000 NB-lncRNAs. Eight clusters were obtained after filtering, normalizing and clustering cells with Seurat (Fig. 3a; L-clusters L0 to L7, where L stands for lncRNA). Based on flow cytometry labels provided in Nguyen *et al* (6), three L-clusters were predominantly luminal and five predominantly basal, characterized by at least 70% of the cells having the corresponding label (Fig. 3a and Table S12). The existence of multiple clusters for each cell type indicated the presence of subpopulations, but as L-clusters were defined based on NB-lncRNA expression only, we could not further characterize them according to protein-coding markers. To circumvent this, we re-clustered the cells based on the expression of GENCODE-annotated genes, characterized the resulting clusters (Fig. 3b; A-clusters, where A stands for annotated) and used this information to label L-clusters. Correspondence between A-clusters and L-clusters was inferred based on the number of cells they shared (Table S12 and Fig. S2).

**Fig. 3.**
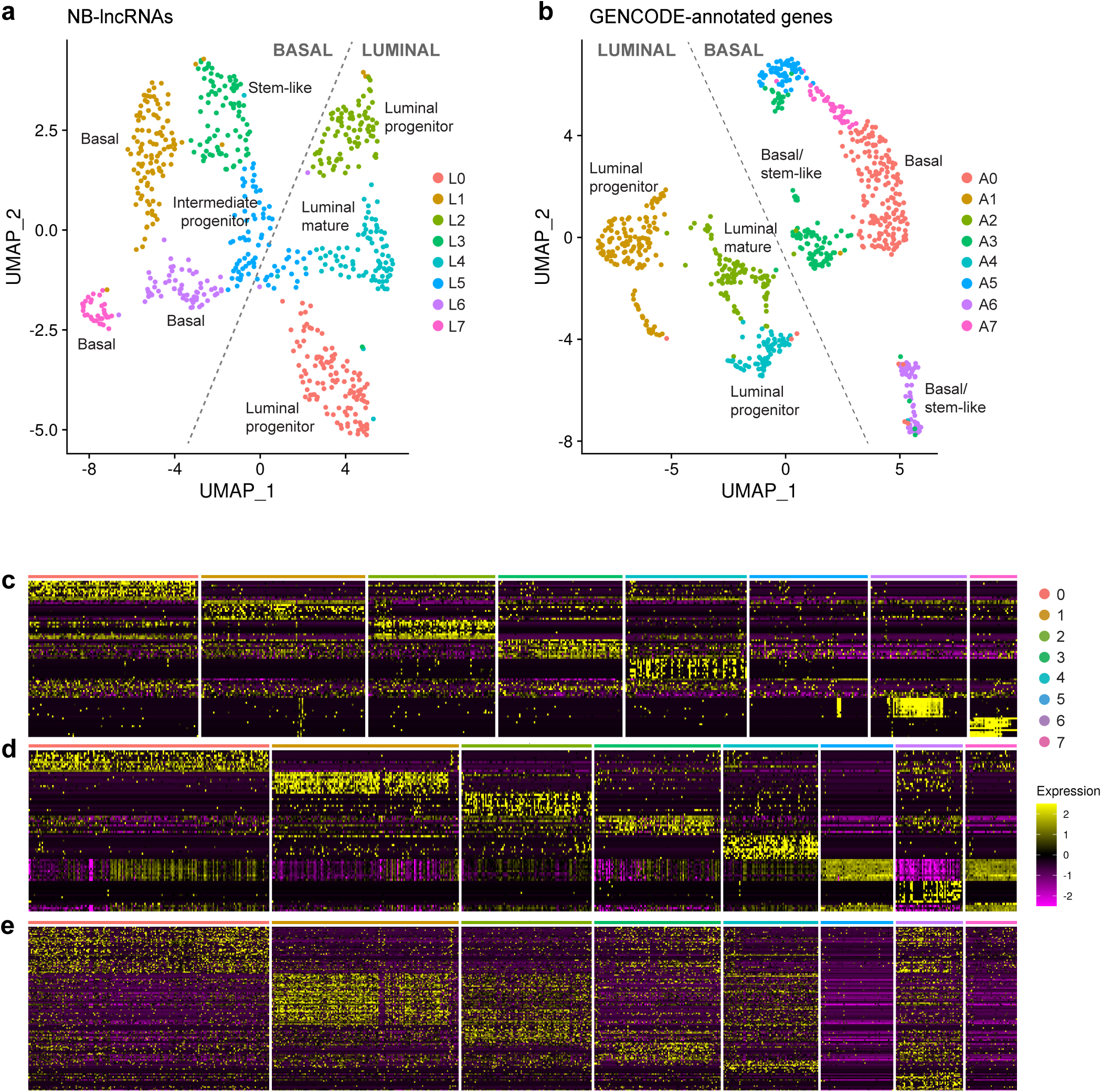
NB-lncRNAs and GENCODE-annotated genes can distinguish breast epithelial cell types. **a** Uniform manifold approximation projection (UMAP) of normal breast cells, clustered based on NB-lncRNAs expression, quantified on Fluidigm scRNAseq data (L-clusters). Cells are color-coded for clusters, which are numbered according to cell counts. **b** UMAP of normal breast cells, clustered based on GENCODE-annotated gene expression, quantified on Fluidigm scRNAseq data (A-clusters). **c** Heatmap showing the top ten Seurat-assigned markers for each L-cluster. **d** Heatmap showing the top ten Seurat-assigned markers for each A-cluster. **e** Heatmap showing Seurat-assigned markers for A-clusters previously reported in the literature.

### Characterization of A-clusters based on known protein markers

Eight A-clusters, also corresponding to three predominantly luminal and five predominantly basal cell populations, were obtained (A0 to A7, Fig. 3b and Table S12). In agreement with Nguyen *et al* (6), the expression of ∼5,000 annotated genes was detected per cell. Seurat uses gene expression patterns to define a set of markers for each cluster with the *FindAllMarkers* function (Table S13). We plotted the top ten most significant Seurat-assigned markers of L-clusters (Fig. 3c) and A-clusters (Fig. 3d) to assess how well they distinguish clusters. For A-clusters, we also plotted markers previously defined in the literature (Table S8 and Fig. 3e) and observed that Seurat-assigned markers define the cell populations better than literature markers.

We first assessed the identity of A-clusters based on the expression of frequently used protein-coding antigens (Table S12 and File S4 (58)). Detection of the main keratin immunohistochemistry marker genes (i.e. *KRT5*, *KRT6* and *KRT14* for myoepithelial and *KRT8* and *KRT18* for luminal cell types) and other relevant protein-coding marker genes (*MME* and *CD44* for basal and *EPCAM*, *MUC1*, *PROM1* and *KIT* for luminal cell types) by Seurat generally coincided with the correspondent FACS labels (Table S12). Physiological characteristics of both luminal and basal cells were also considered. For example, the sets of Seurat-assigned markers of clusters A0 and A6 were the most significantly enriched for EMT-related genes and genes characteristic of claudin-low status (enrichment p-values < 4.0 e-35; Tables S8 and S12), two hallmarks of basal cells (59–61). Cells in the luminal compartment are known to have shortened telomeres, which elicit DNA repair (62). Markers of clusters A2 and A4 were the most significantly enriched on genes involved in telomere maintenance and markers of cluster A4 on DNA repair genes (enrichment p-values < 8.0e-03; Tables S8 and S12).

To differentiate luminal progenitors from luminal mature subpopulations, we used the annotated gene markers obtained by Pal et al (56) based on scRNAseq of normal mammary glands. When compared to Seurat-assigned markers of each A-cluster, these markers unequivocally (enrichment p-values < 1.0e-10) characterized clusters A0, A3 and A6 as basal, clusters A1 and A4 as luminal progenitor and cluster A2 as luminal mature (Table S12). Both luminal progenitor clusters (A1 and A4) also express higher levels of *H2B* genes, shown to have short-term repopulating potential in mice (63). In addition, progenitor cluster A1 is associated with highest expression of luminal alveolar and hormone-sensing markers (e.g. *EHF*, S100A6), whereas A4 has high expression of KIT, which is a hallmark of ductal progenitors and cluster A2 has a classical mature ductal cell signature (FOXA1^hi^ and ELF5^low^) and hormone-sensing characteristics (highest expression of *AREG*, *CITED1*, *LY6D* and *PRLR*) (64–66). Notably, all three luminal clusters are devoid of *ESR1* and *PGR* expression.

We completed the annotation of A-clusters comparing Seurat-assigned markers with an in-house dataset comprising 359 markers from the literature (Tables S12 and S8, File S2 and Fig. 3e). Based on the overlap between these gene lists, we confirmed cluster A0 as basal with strong myoepithelial signature (45% of the known markers), clusters A1 and A4 as luminal progenitor (25% and 28% of the known markers) and cluster A2 as luminal mature (13% of the known markers). Clusters A3 and A6 have basal origins and the highest number of breast stem-cell markers identified by Seurat (12 and 16 known markers respectively, or 37% and 31% with enrichment p-values < 1.0e-10). Markers in both clusters include *JAG1*, *ITGB1* and *YBX1*, while cluster A3 also expresses high levels of *CD44*, *ZEB1* and others and cluster A6 of GNG11, TCF4 and others. Clusters A5 and A7 remained undefined, with no known markers in the list generated by Seurat. Additional investigation showed evidence of cell death occurring in both of these clusters (based on mitochondrial gene markers, File S4) and traced cells to basal origins (correspondence p-values 0.01 and 2.0e-11, respectively).

### Further characterization of subpopulations represented by L-clusters

We labeled each cell according to the A-cluster they were assigned to and characterized L-clusters based on their cell composition (Fig. S2 and Table S12). More than 95% of the cells in cluster L0 are in luminal progenitor cluster A1. Cluster L1 is predominantly basal, with ∼90% of its cells in cluster A0. Cluster L2 has ∼80% of its cells in cluster A4 (luminal progenitor) and ∼15% in cluster A2 (luminal mature). Cluster L3 represents a heterogeneous progenitor population, with ∼65% of its cells in cluster A6 and ∼30% in cluster A3 (both of which have high numbers of stem-cell markers). Cluster L4 has ∼90% of its cells in cluster A2 (luminal mature). Cluster L5 is a mixed population with comparable portions of cells in clusters A1 (∼15%), A3 (∼25%), A6 (∼15%) and A7 (∼20%). Clusters L6 and L7 are mainly (respectively ∼70% and ∼90%) formed by cells of cluster A0 (basal). In summary, luminal clusters obtained based on GENCODE-annotated gene expression were replicated using NB-lncRNAs expression, while the heterogeneous basal cluster A0 is subdivided into three clusters L1, L6 and L7. Doubling Seurat’s resolution parameter for A-clusters subdivided cluster A0 in two cell subpopulations, one mainly (>80%) represented by cluster L1 and another by clusters L6 and L7. The subpopulation that forms cluster L1 is marked by the expression of myoepithelial genes, such as hemidesmosome components (*COL17A1*, *LAMA3* and *LAMC1*) and actin-binding genes (*CALD1*, *MYLK* and *SVIL*). The other subpopulation had more diverse markers, including genes with roles in vascularization and innervation (e.g. F3, PDGFA, SFRP1, SOD2 and TNC) and wound healing (*CAV1* and *PLAU*), which are characteristic of the stroma. Cluster L5 is a mixed cluster which contains cells from different compartments and may represent a subpopulation of intermediate progenitor cells. Stem-like clusters A3 and A6 are merged into cluster L3, which congregates nearly twice as many stem-cell marker genes than any other cluster.

Using Seurat we identified many NB-lncRNAs as potential biomarkers (Tables S13 and S14). For each L-cluster, ∼15% of all NB-lncRNA markers are intergenic, ∼30% intronic and ∼8% antisense. Examples of luminal markers are: (i) DN86902C0G3I1 (cluster L0), transcribed from ERO1A, a gene that promotes metastatic colonization in breast cancer (67); (ii) DN574858C0G1I1 (cluster L2) antisense to the *STAT3* activator *SIX4* and (iii) DN91892C0G2I1 (cluster L4), intronic to ZEB1 (Fig. 4a). Examples of basal markers are (iv) DN690057C0G1I1 (cluster L1), intergenic to breast progenitor marker YBX1; (v) DN21900C0G1I1 (cluster L6), intergenic to quiescence-regulator *SATB2* and (vi) DN87246C0G1I6/*FOXG1-AS1* (cluster L7), antisense transcript of proliferation-related gene FOXG1 (Fig. 4b). Examples of NB-lncRNAs that regulate or are co-regulated with protein-coding genes are general basal marker *NCF4-AS1* (DN1070C0G3I1; clusters L1, L6, L7), and general luminal marker *AC005077.7* (DN262C0G1I12; clusters L0, L2 and L4). Both NB-lncRNAs mirror the expression of their protein-coding counterparts (Figs. 4c, d), suggesting functional relationships.

**Fig. 4.**
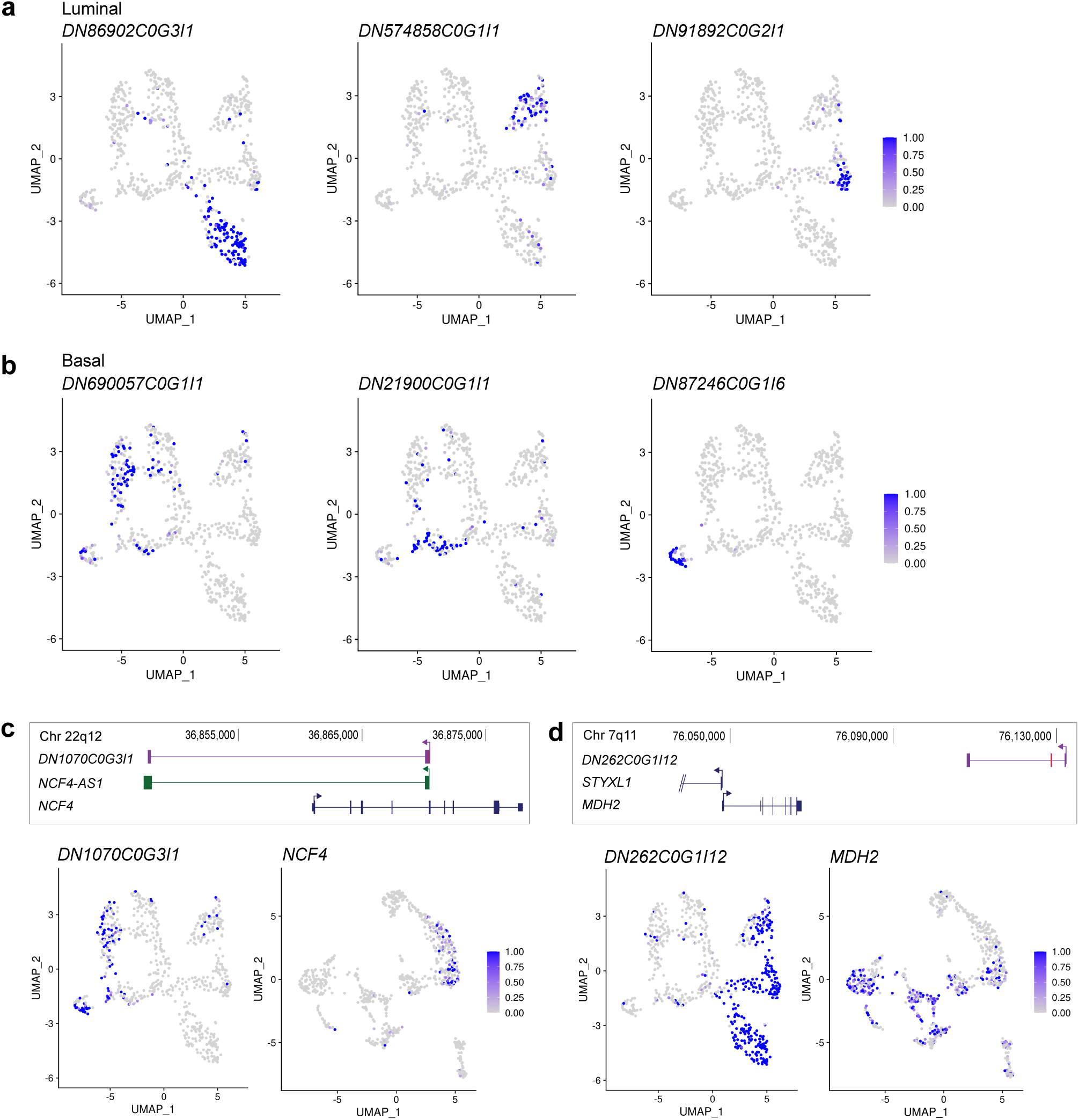
Seurat-assigned markers identified candidate NB-lncRNA biomarkers. **a** Expression patterns of three identified NB-lncRNA luminal markers in L-clusters. **b** Expression patterns of three identified NB-lncRNA basal markers in L-clusters. **c, d** Upper panels: UCSC genome browser (hg38) diagrams showing NB-lncRNAs (purple), GENCODE-annotated protein-coding (blue) or noncoding genes (green). Lower panels: Expression patterns of NB-lncRNAs in L-clusters (left) and correlated protein-coding genes in A-clusters (right), showing examples of co-expression between coding and noncoding genes at single-cell level.

Importantly, clustering of cells acknowledging NB-lncRNA expression (i.e. L-clusters and M-clusters) seems to result in a robust and accurate representation of breast epithelial cell populations. For example, NB-lncRNAs perform better than annotated lncRNAs for grouping cells with similar FACS labels in the same clusters. Overall, based on confusion matrices, M-clusters perform the best at resolving the cell types (Table 1). Additionally, 69% of the Seurat-defined NB-lncRNAs for L-clusters are cluster-specific, which is another measure of cluster quality. For comparison, only 63% of the Seurat-defined annotated lncRNA markers were cluster-specific. Finally, clustering trees reveal L-clusters to be stable, with no evidence of cluster instability or of unexpected cell mobility between clusters as resolution is increased (Fig. S3).

**Table 1.** Measures of cluster performance based on confusion matrices of correctly and incorrectly classified cells, according to available FACS labels. The F1 score is defined as 2xTP/(2xTP + FP + FN). The Matthews Correlation Coefficient (MCC) is defined as TP*TN - FP*FN / sqrt((TP+FP)*(TP+FN)*(TN+FP)*(TN+FN)). TP: true positives, FP: false positives, TN: true negatives and FN: false negatives.

| Metric | NB-lncRNAs | GENCODE lncRNAs | GENCODE | NB-lncRNAs +<br>GENCODE |
| --- | --- | --- | --- | --- |
| True positives | 346 | 396 | 409 | 412 |
| False positives | 26 | 43 | 15 | 13 |
| True negatives | 267 | 255 | 286 | 289 |
| False negatives | 16 | 16 | 27 | 27 |
| Sensitivity | 0.9558 | **0.9612** | 0.9381 | 0.9385 |
| Specificity | 0.9113 | 0.8557 | 0.9502 | **0.9570** |
| Precision | 0.9301 | 0.9021 | 0.9646 | **0.9694** |
| Accuracy | 0.9359 | 0.9169 | 0.9430 | **0.9460** |
| F1 score | 0.9428 | 0.9307 | 0.9512 | **0.9537** |
| MCC | 0.8703 | 0.8298 | 0.8833 | **0.8897** |
| FDR | 0.0699 | 0.0979 | 0.0354 | **0.0306** |

MCC: Matthews Correlation Coefficient
FDR: False Discovery Rate

### LncRNA expression better defines breast subpopulations than protein-coding genes

We compared the expression pattern of lncRNAs and protein-coding genes at single-cell level. To minimize bias, we simultaneously quantified protein-coding genes, annotated lncRNAs and NB-lncRNAs. First, we analyzed broadly expressed genes (defined as detectable expression in >1/3 of the cells) or restricted (<1/3 of the cells) expression patterns and investigated differences between NB-lncRNAs and protein-coding genes. More than 25% of the protein-coding genes were broadly expressed in normal breast cells, but only 6% of the annotated lncRNAs and <1% of the NB-lncRNAs (Fig. 5a, Table S15 and File S4). On average, protein-coding genes were expressed in ∼140 cells, while annotated lncRNAs were expressed in ∼30 and NB-lncRNAs in ∼15 (Fig. 5b). In agreement with their expected biological function, the set of broadly expressed protein-coding genes is enriched in known housekeeping genes (47%, p-value ∼ 0; Table S15). Accordingly, protein-coding genes nearest or co-expressed with the 121 broadly expressed NB-lncRNAs were also enriched for known housekeeping genes (46% with p-value 5.4e-31 and 35% with p-value 9.6e-19; Table S15) e.g. *GAPDH*, *HSPA5*, *IAH1*, *MKLN1*, *PSMD7*, *SYNCRIP* and *TIMM44*. Analysis of genes with restricted expression patterns showed the median cellular expression of NB-lncRNAs was nearly twice that of protein-coding genes (p-value < 1.9e-06; Fig. 5c).

**Fig. 5.**
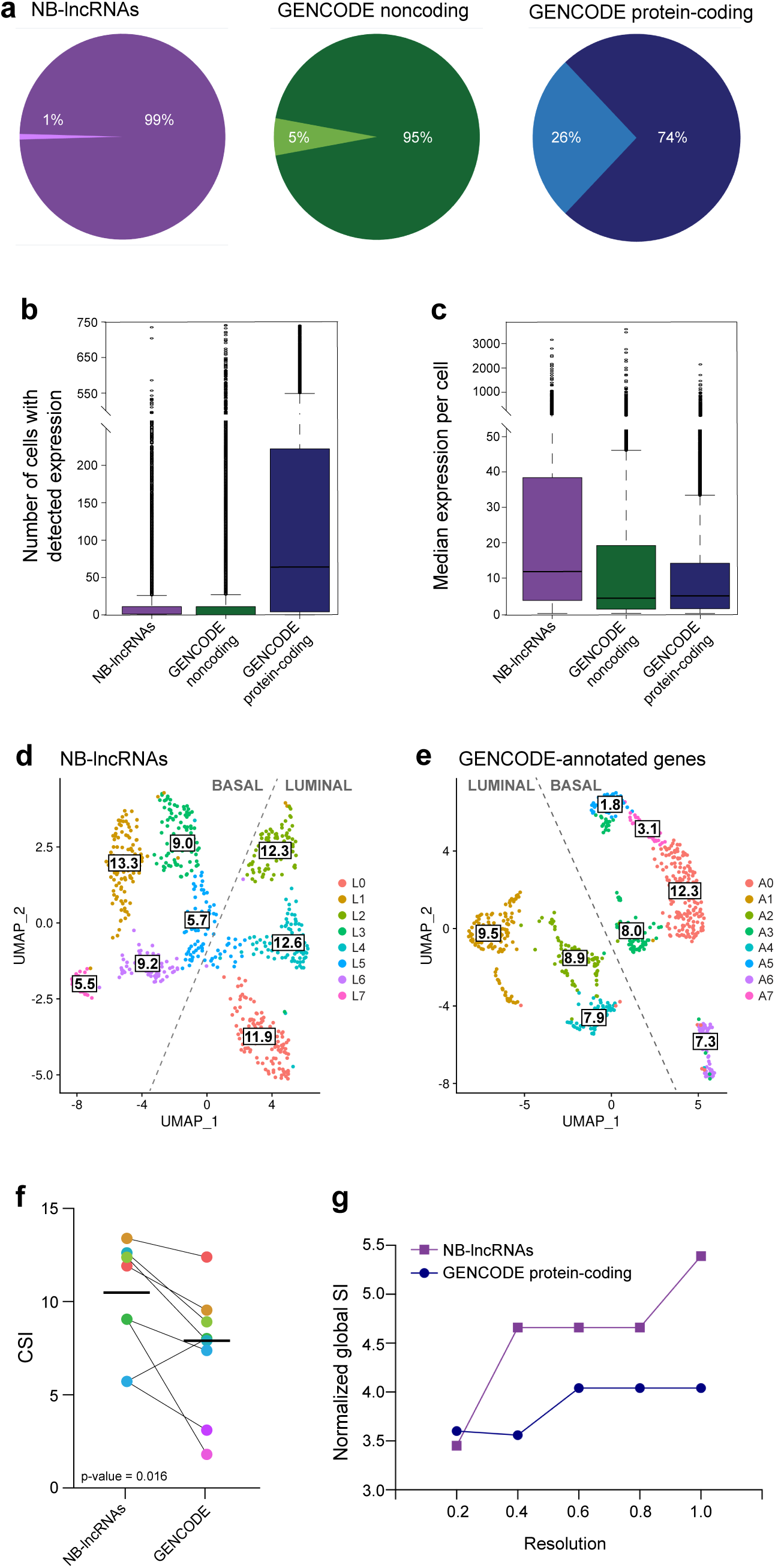
NB-lncRNAs have compartmentalized expression levels, which are higher at cell-level. **a** Proportion of NB-lncRNAs (purple) or GENCODE-annotated noncoding (green) or protein-coding (blue) genes with restricted (darker) versus widespread (lighter) expression patterns. **b** Boxplot of the number of cells in which NB-lncRNAs (purple) or GENCODE-annotated noncoding (green) or protein-coding (blue) genes are expressed, from the total 741 cells. On average, NB-lncRNAs are expressed in ∼15 cells, GENCODE-annotated noncoding genes in ∼30 cells and GENCODE-annotated protein-coding genes in ∼140 cells. **c** Boxplots of the median expression of NB-lncRNAs (purple) or GENCODE-annotated noncoding (green) or protein-coding (blue) genes per cell (in TPMs), showing NB-lncRNAs have higher cell-level expression. **d** UMAP showing normal breast cell clusters obtained based on NB-lncRNA gene expression (L-clusters), with their corresponding cluster specificity index (CSI). **e** UMAP showing normal breast cell clusters obtained based on GENCODE-annotated gene expression (A-clusters), with their corresponding CSI. **f** Dotplots showing the difference in CSI for corresponding clusters in ‘**d’** and ‘**e’**. Dots were colored according to the represented L-cluster or A-cluster, bold horizontal lines mark the average CSI for each gene set and the p-value (0.016; Fisher’s exact test) shows the difference was significant. **g** Increase in normalized global SI levels, obtained as the normalized average of the CSIs of all clusters, as resolution is increased.

Since NB-lncRNAs have a more restricted expression, we expected them to better define breast epithelial subpopulations than protein-coding genes. To test this, we first calculated the specificity index (SI) of each transcript in each cluster, which is their median expression in each cell of that cluster divided by the sum of their median expression in all clusters (based on (37)). We then defined the cluster specificity index (CSI) as the average SI of all transcripts in each cluster. CSIs for L-clusters were always higher than for the corresponding A-cluster (Figs. 5d-f; p-value 0.016), indicating that NB-lncRNA expression is more subpopulation-specific. For comparison, we re-clustered cells based on the expression of ∼23,000 GENCODE-annotated genes that are neither protein-coding or confirmed lncRNAs (referred to as O-clusters, where O stands for other noncoding transcripts) and observed an overall decrease in CSI of correspondent clusters (Fig. S4; p-value 0.4). This confirmed that higher cluster specificity is a characteristic of NB-lncRNAs.

We reasoned that, having higher cluster specificity could be a reflection of restricted NB-lncRNAs having compartmentalized expression within clusters, being limited to a subset of cells. To assess this, we enforced overclustering of normal breast cells, by increasing the resolution parameter in the *FindClusters* function of Seurat. Consistent with our hypothesis, increasing the number of clusters increased the specificity index of NB-lncRNAs but not of protein-coding genes. Indeed, gradually varying the resolution from 0.2 to 1.0 resulted in an 59% global increase in the global normalized SI of NB-lncRNAs, while for protein-coding genes the increase was of only 11% (Fig. 5g). Additionally, 30% of the NB-lncRNAs but only 13% of the protein-coding genes became more cluster-specific (i.e. had higher global normalized SI) as resolution was increased.

### NB-lncRNAs are highly tissue-specific

As the expression patterns of NB-lncRNAs confirmed their high cell subpopulation-specificity, we reasoned they would also be tissue-specific. To assess this, we used data from a recent publication (34), where authors assessed the transcriptomes of several normal human tissues. An in-house RNAseq analysis pipeline was used to estimate the abundance of individual transcripts in 27 tissues. Tissues were ranked according to the occurrences of highest tissue-specificity in different transcript subclasses. This analysis revealed NB-lncRNAs were highly breast-specific, as opposed to GENCODE-annotated genes (either protein-coding or lncRNAs), which were not specifically enriched in the breast tissue (Fig. 6a). We further listed the top 1,000 most breast-specific NB-lncRNAs and investigated their expression in all 27 tissues (Fig. S5), confirming their tissue-specificity.

**Fig. 6.**
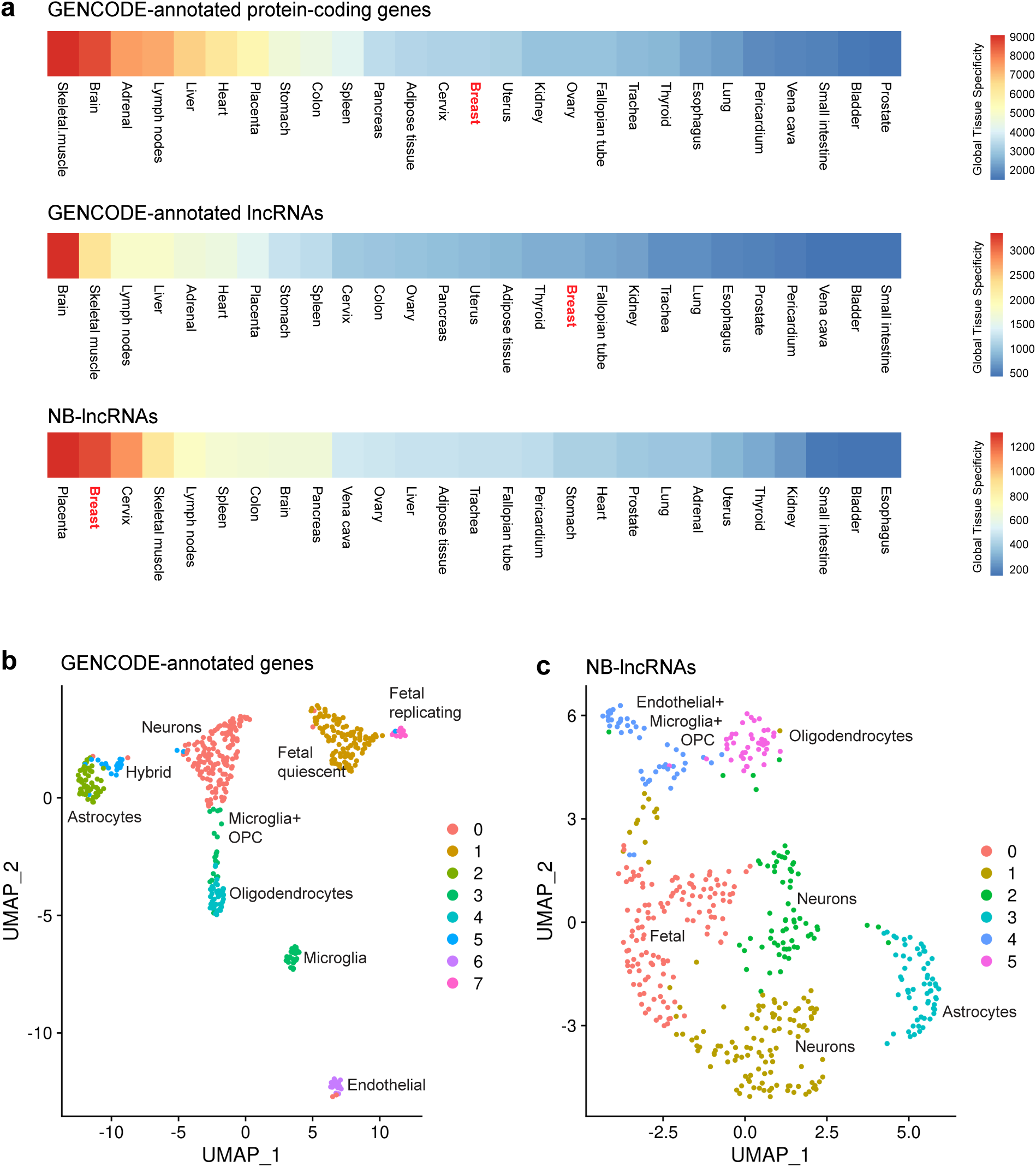
NB-lncRNAs cannot identify human brain cell types. **a** Global tissue-specificity of annotated protein-coding genes (top), annotated lncRNAs (middle) and NB-lncRNAs (bottom) across 27 human tissues. The tissue-specificity was summarized by calculating the tau index of every transcript in each set and counting the number of occurrences where each tissue had the highest tau overall. **b** UMAP of normal breast cells, clustered based on GENCODE-annotated gene expression, quantified on Fluidigm scRNAseq data. Cells are color-coded for clusters, which are numbered according to cell counts. **c** UMAP of normal breast cells, clustered based on NB-lncRNAs expression, quantified on Fluidigm scRNAseq data. Clusters were labeled based on the information provided in (35) and known protein-coding markers of the represented brain cell types.

### NB-lncRNAs poorly define cell populations in the human brain

We reasoned that, being highly tissue-specific, NB-lncRNAs would not perform well at defining cell populations in other tissues. We tested their ability to cluster brain cells using Fluidigm scRNAseq data derived for 466 cells isolated from healthy human brain samples (35) (File S2). Cells were sequenced at nearly twice the average depth of the normal breast cells (∼3M versus ∼1.6M reads). We first clustered cells based on the expression of GENCODE-annotated genes, obtaining clusters consistent with those in Darmanis *et al* (35) (Fig. 6b). Notably, we retrieved several important markers of immunoreactive microglia (e.g. *CD74*, *CCL4*, *AIF-1*, *ALOX5AP*), cortical cells (e.g. *HSF2*, *NEUROD6*) and brain development (e.g. *IGFBPL1*, *FABP7*, *LMO3*). We then used the expression of NB-lncRNAs for cell clustering. On average, ∼1,100 NB-lncRNAs were expressed per brain cell, but this did not result in a successful clustering of the different brain cell subpopulations (Fig. 6c). In fact, although NB-lncRNAs expression could coarsely group cell types, the clustering failed to unambiguously discern subpopulations of oligodendrocyte precursor cells (OPCs), microglia and endothelial cells and to distinguish fetal quiescent from fetal replicating cells (Table S16). Overall, clustering based on NB-lncRNAs expression performed poorly compared with clustering based on GENCODE-annotated genes expression, despite considerably higher sequencing depth. This supports our claims that lncRNA discovery should routinely be performed on similar samples a priori, to improve scRNAseq clustering.

### NB-lncRNA gene expression defines a subpopulation of breast stem-cells

We used three methods to ascertain which L-cluster most likely harbors breast stem-cells. (i) Protein-coding gene expression at the cell level was used to calculate the average expression of putative stem-cell markers in human and mouse mammary tissue (*ATXN1*, *BMI1*, *CD1D*, *GNG11*, *ID4*, *INPP5D*, *ITGB1*, *ITGB3*, *JAG1*, *MME*, *PARD3B*, *PROCR*, *TCF4* and *ZEB1*; Table S17) per L-cluster (Fig. 7a). (ii) We assessed the expression of genes involved in the three main pathways associated with stemness in a number of tissues: Notch, Hedgehog and Wnt signaling (Fig. S6). (iii) The Single-Cell ENTropy (SCENT) method (38) was used to estimate the differentiation potency of each cell, based on how efficiently signaling can diffuse through a gene interaction network given the cell gene expression profile. Conceptually, cells with higher differentiation potency express more connected genes, which results in higher entropy (i.e. the SR value in SCENT). We modified the gene interaction network of SCENT including annotated lncRNAs based on correlation data obtained from LncRNAs2Pathways (39). SR values were calculated using cell-level annotated gene expression averaged across L-clusters (Fig. 7b and Table S17). All three methods confirmed cluster L3 as having the highest degree of stemness. Cluster L3 is a mix of ∼65% cluster A6 and ∼30% cluster A3 (both of which were shown to express high levels of stem-cell markers during A-cluster characterization; Fig. 3b).

**Fig. 7.**
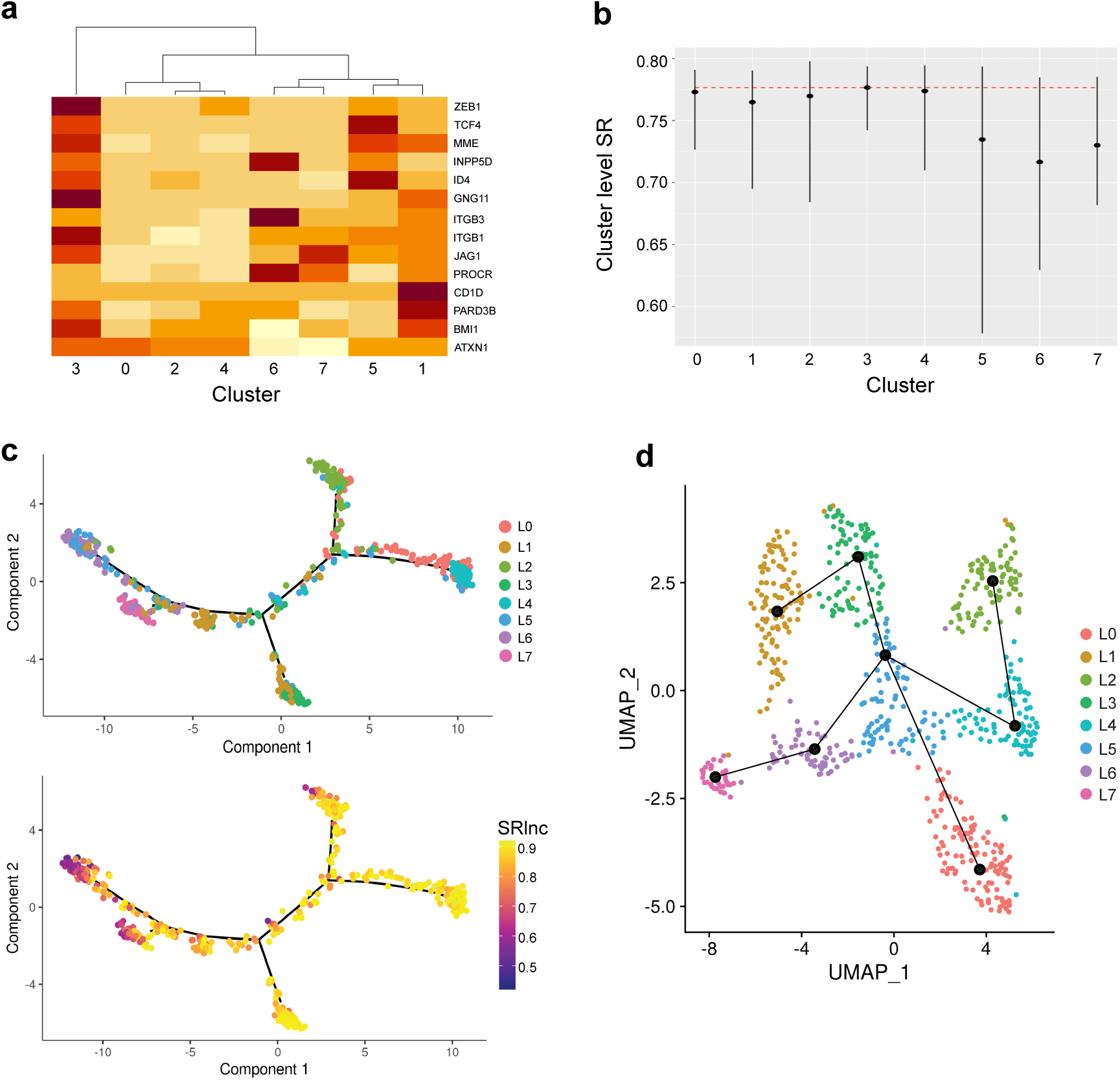
NB-lncRNAs define a stem-like cell subpopulation in the normal breast epithelium. **a** Heatmap of the average expression of genes with experimentally confirmed stem-cell properties across the L-clusters. Genes were selected for their reported capacity of repopulating mammary fat pads, originating both basal and luminal compartments. Expression was measured at cell level and averaged across cells allocated to the same L-cluster. **b** The SCENT method was used to calculate the signaling entropy level (SR value) of each cell. Cells in the same L-cluster were plotted together, with the average for each cluster represented by dots. The dashed red line was placed at the highest average (cluster L3). **c** Monocle plots with cells distributed along a differentiation trajectory, with branch points representing cell lineages. Cells were colored either by Seurat-assigned L-clusters (upper) or by SR value (lower). **d** Slingshot trajectories showing the placement of lineages on top of the UMAP of L-clusters obtained with Seurat. Cluster L3 was defined as the root state, based on its higher stemness.

We used Monocle to visualize the diffusion of signaling entropy along the mammary epithelial cell hierarchy (Fig. 7c). The placement of clusters and branch points broadly confirmed cell lineages. The branch containing L3, defined as root, which radiates through basal cluster L1 to a bough of more differentiated basal clusters (L5 and L6 in one arm and L7 on a separate arm) and another bough which bifurcates into a luminal progenitor cluster L2 on one side and luminal clusters L0 and L4 on the other side. The placement of clusters L0 and L4 in the same branch may reflect their observed higher proliferative rates, hormone-sensitivity or other characteristics. Being confident that the distribution of SR values along the cell hierarchy recapitulated the mammary differentiation process, we used Slingshot to infer corresponding trajectories, defining cluster L3 as the root state (Fig. 7d). Consistent with other studies (6,68), the results confirmed the existence of a cell population with characteristics of a common progenitor for basal and luminal cell types. The cell hierarchy suggests differences between the basal L-clusters which subdivided from cluster A0, with clusters L1 closer to L3 and clusters L6 and L7 to L5, which appears to act as an intermediate progenitor. Indeed, cluster L5 is formed by cells with diverse gene expression profiles and markers of both basal and luminal cell types. Cluster L5 is also a precursor to luminal clusters L0 on one branch and L2 and L4 on another, confirming observations made with Monocle.

### NB-lncRNAs also define breast cell subpopulations in 10x Genomics scRNAseq data

As the 10x Genomics platform has surpassed Fluidigm as the method of choice for scRNAseq, we wanted to ensure our results were transferable and not dependent on the higher depth and sequencing strategy of the Fluidigm platform. Using CellRanger, we clustered 10x Genomics scRNAseq data from four individuals (also obtained from Nguyen *et al* (6)) into cell subpopulations (Figs. 8a-c for individual number 4 and Fig. S7 for individuals 5 to 7). Individual number 4 was studied in more detail. On average, we detected the expression of >200 NB-lncRNAs and >1,800 annotated genes per cell. NB-lncRNA expression alone has successfully segregated cells of all individuals into subpopulations. Seven, ten and twelve clusters were respectively obtained based on either NB-lncRNA expression, GENCODE-annotated gene expression or a merge of both gene sets (Figs. 8a-c). Specific markers were found for each cell subpopulation (Fig. S8a). We followed the same strategy used with the Fluidigm data to characterize clusters based on known protein-coding markers (Table S8) and define cluster correspondence. The cell type-dependent structure was broadly retained between clustering experiments. Differences were observed in the number of clusters characterized by breast stem markers (Figs. 8a-c, Fig. S8b), including CD44, GNG11, TCF4 and YBX1. Notably, these clusters were not marked by keratin genes and their epithelial origin could not be confirmed.

**Fig. 8.**
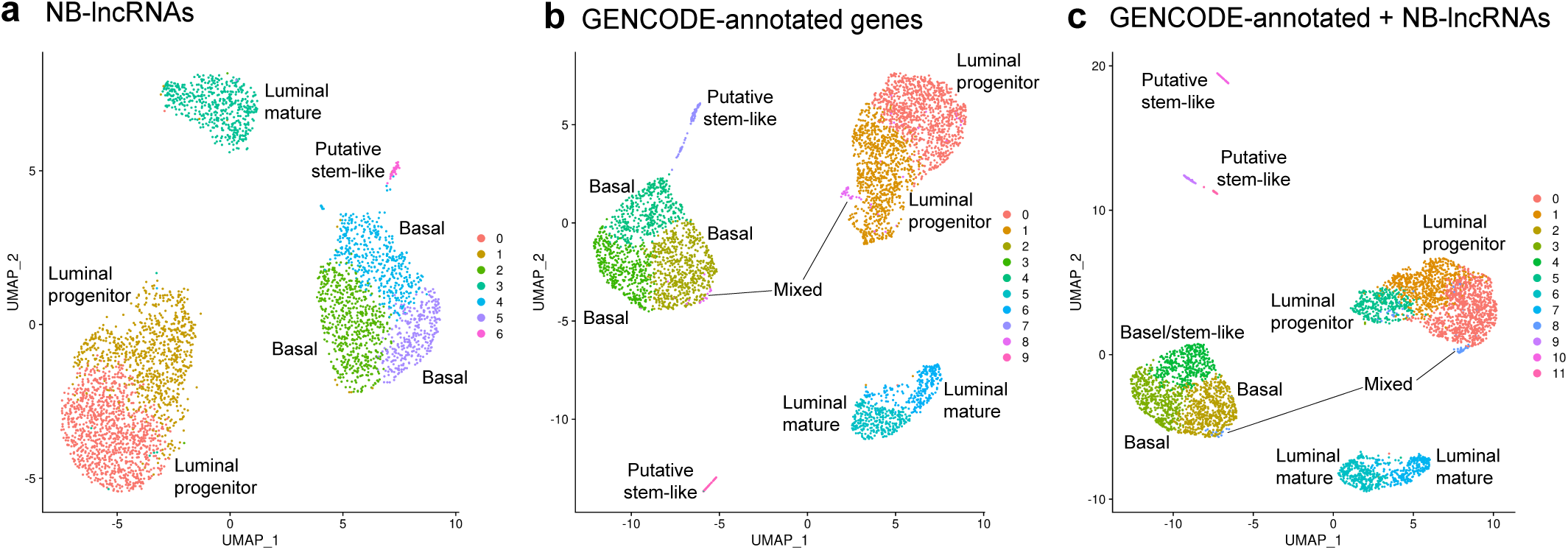
NB-lncRNAs discern normal breast cell subpopulations on 10x Genomics scRNAseq. **a** UMAP of normal breast cells, clustered based on NB-lncRNAs expression, quantified on 10x Genomics scRNAseq data. Cells are color-coded for clusters, which are numbered according to cell counts. **b** UMAP of normal breast cells, clustered based on GENCODE-annotated gene expression, quantified on 10x Genomics scRNAseq data. Cluster labels are based on the presence of known markers of cell subpopulations in the list of Seurat-assigned markers. **c** UMAP of normal breast cells, clustered based on the expression of GENCODE-annotated genes and NB-lncRNAs, quantified on 10x Genomics scRNAseq data.

While clustering based on GENCODE-annotated genes resulted in two putative stem-like clusters, combining their expression with that of NB-lncRNAs resulted in three. Moreover, the luminal progenitor population was further subdivided, creating another additional cluster. To investigate how NB-lncRNA expression could be contributing to the emergence of the additional clusters, we re-clustered cells based on GENCODE-annotated genes alone, increasing the resolution parameter in Seurat. While forced overclustering also resulted in the subdivision of the luminal population into three clusters, no additional putative stem-like clusters were observed (Fig. S8c). This suggested NB-lncRNAs might be involved in stemness and in differentiating stem-cell states in the normal human breast.

### Correlations between NB-lncRNAs and breast cancer

To establish links between NB-lncRNAs and breast cancer, we assessed their expression levels in tumor samples of different molecular subtypes from The Cancer Genome Atlas (TCGA (42) with subtypes defined as per (43)). TCGA RNAseq data was remapped to the human transcriptome supplemented with the NB-lncRNAs and gene expression was re-quantified. Significant correlations (Pearson correlation p-values < 0.05) were observed between NB-lncRNA expression profiles in TCGA subtypes and L-clusters, especially luminal cluster L4 and basal cluster L1 (Table S18). Furthermore, expression of Seurat-assigned NB-lncRNA markers for L-clusters successfully stratified TCGA samples into basal and luminal molecular subtypes (Fig. 9a) to a greater extent than Seurat-assigned markers of A-clusters (Fig. 9b). This indicated a relevant contribution of NB-lncRNA gene expression to the molecular classification of breast tumor subtypes.

**Fig. 9.**
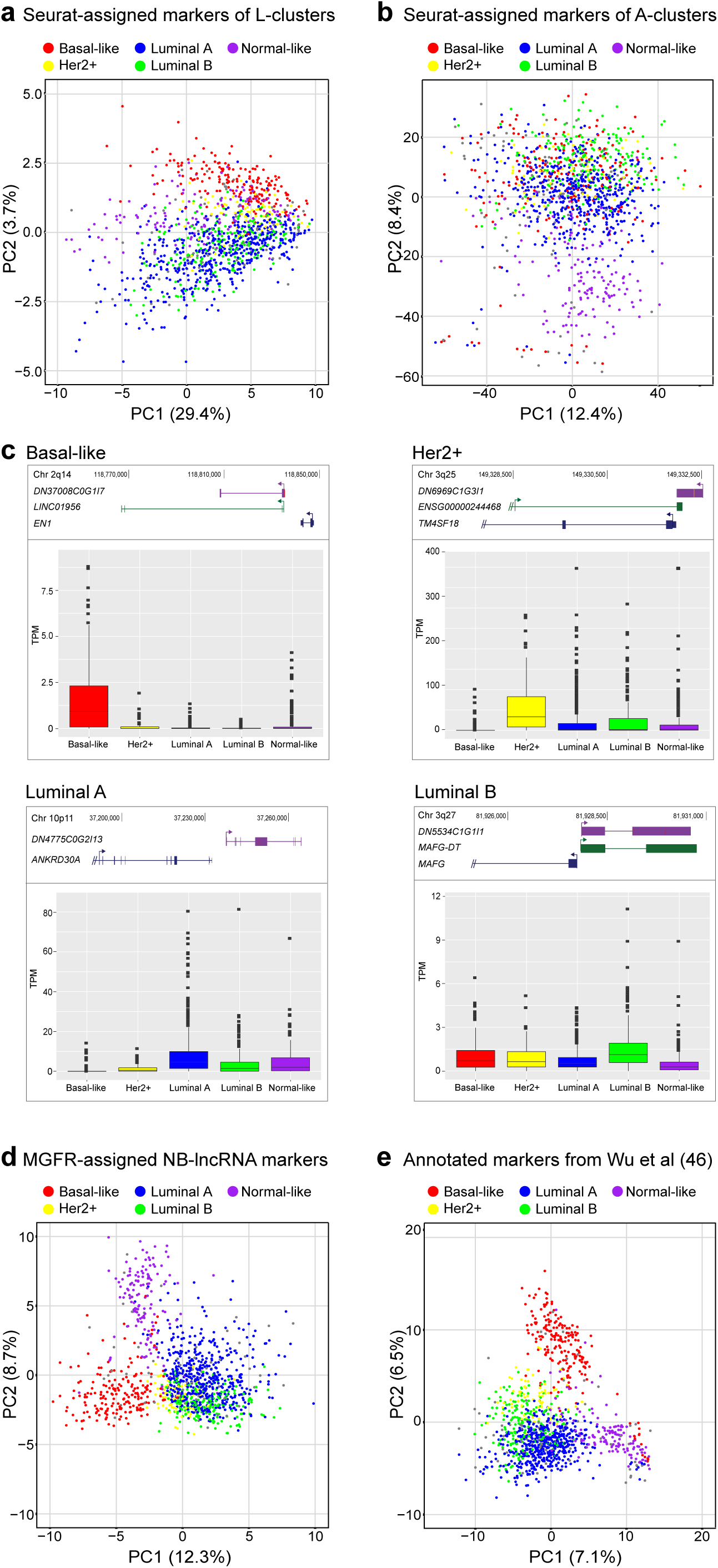
NB-lncRNA expression can differentiate between breast cancer subtypes in TCGA. **a** The expression of Seurat-assigned NB-lncRNA markers of L-clusters separates the main subtypes of TCGA breast cancer tumors in a two-dimension PCA plot. **b** To a lesser degree, the expression of Seurat-assigned GENCODE-annotated markers of A-clusters separates different subtypes of TCGA breast cancer tumors in a two-dimension PCA plot. **c** NB-lncRNAs are markers of specific breast cancer subtypes. Four selected NB-lncRNAs are shown with their genomic context (upper; UCSC genome browser diagram) and expression levels (lower; boxplots of TPMs in TCGA samples of each subtype). **d** In-house NB-lncRNA markers of TCGA breast cancer subtypes can separate all subtypes of breast cancer in a two-dimensional PCA plot. **e** For comparison, known breast cancer protein-coding markers separate subtypes with comparable performance.

To further explore the correlation of NB-lncRNAs with breast cancer subtypes, we derived lists of NB-lncRNA markers for each TCGA breast cancer subtype, using an in-house method based on MGFR (45), with bootstrapping. Approximately 80% of the selected NB-lncRNA markers (Table S18) are novel genes or novel isoforms of annotated lncRNA genes. Notably, among the identified NB-lncRNA markers there were 28 experimentally-supported lncRNAs with established correlations with breast cancer (53–55) (Table S9; overlap p-value < 0.001), including *LINC01089*, *MEG3*, *HOXB-AS1*, *HOXC-AS1* and *TTC39A-AS1* (Table 2; which also contains known breast cancer-related lncRNAs found as markers of L-clusters). There were also many novel NB-lncRNAs identified as subtype-specific (e.g. Fig. 9c; Table S18). For example, MGFR-assigned Her2^+^ subtype marker DN6969C1G3I1 overlaps an annotated gene (ENSG00000244468) previously listed as one of the most highly expressed lncRNAs in breast cancer (69).

**Table 2.** Annotated NB-lncRNAs markers of normal breast cell subpopulations (top) or TCGA breast cancer subtypes (bottom). Top: Seurat-assigned NB-lncRNA markers with previous annotation, experimental evidence and established links with breast cancer are listed with the corresponding normal breast cell subpopulation for which they are markers and short identifiers. Bottom: MGFR-assigned NB-lncRNA markers with previous annotation, experimental evidence and established links with breast cancer are listed with the corresponding breast cancer subtype for which they are markers and short identifiers. Full identifiers are available in Table S18.

| Annotated lncRNA | Cell subpopulation(s) | NB-lncRNA(s) |
| --- | --- | --- |

| *CARMN* | General basal | DN405 |
| --- | --- | --- |
| *LINC01094* | Basal | DN19901 |
| *MEG3* | Basal | DN4816 |
| *MAGI2-AS3* | Basal | DN935 |
| *HOTAIRM1* | Basal | DN4816, DN126378 |
| *SNHG29* | General luminal | DN1842 |
| *DANCR* | Luminal mature | DN1852 |
| *ZFHX4-AS1* | Luminal mature | DN53411 |
| *LINC00993* | Luminal mature | DN4775 |
| *LINC01151* | Luminal | DN20235 |
| *LINC01089* | Luminal progenitor | DN11934 |
| *SNHG15* | Stem-like | DN141 |
| *GAS5* | Stem-like, intermediate progenitor | DN798 |
| Annotated lncRNA | TCGA subtype(s) | NB-lncRNA(s) |
| *FOXP4-AS1* | Basal | DN8672 |
| *RP11-132A1.4* | Basal | DN98035 |
| *SBF2-AS1* | Basal, Her2<sup>+</sup> | DN13758 |
| *HOXC-AS1* | Her2<sup>+</sup> | DN30804 |
| *LINC01133* | Her2<sup>+</sup> | DN3404, DN364571 |
| *MNX1-AS1* | Her2<sup>+</sup> | DN87774 |
| *OIP5-AS1* | Her2<sup>+</sup> | DN167605, DN733 |
| *RP11-206M11.7* | Her2<sup>+</sup> | DN6969 |
| *RP11-390F4.3* | Her2<sup>+</sup> | DN64622 |
| *ST8SIA6-AS1* | Her2<sup>+</sup> | DN68716 |
| *EPB41L4A-AS1* | Luminal A | DN17927 |
| *HOXB-AS3* | Luminal A | DN12537 |
| *LINC00993* | Luminal A | DN839C |
| *PCAT18* | Luminal A | DN38481 |
| *TTC39A-AS1* | Luminal A, Luminal B | DN58767 |
| *LINC01089* | Luminal B | DN11934 |
| *MAFG-AS1* | Luminal B | DN5534 |
| *HOTAIRM1* | Normal-like | DN126378 |
| *MEG3* | Normal-like | DN4816 |
| *SNHG18* | Normal-like | DN14445 |

To assess the performance of our strategy, we used the same method to generate protein-coding gene marker lists for each subtype and compared these to markers obtained by Wu *et al* (46), confirming significant correspondence (p-values of overlaps < 0.002; Table S18). We then assessed the correspondence between the MGFR-assigned NB-lncRNAs markers and protein-coding markers of TCGA breast cancer subtypes (46), by listing all annotated genes within 500kb of each NB-lncRNA marker and overlapping these with the markers from Wu *et al* (Table S18). With the exception of the luminal B subtype, the sets of annotated genes nearby NB-lncRNA markers were enriched (overlap p-values < 0.02) in reported protein-coding markers of the corresponding breast cancer subtype (46). Genes nearby NB-lncRNA markers of the luminal B subtype were enriched in reported luminal A markers, likely reflecting the similarities between these two subtypes. Indeed, there was a significant overlap (p-value < 0.01) between genes near MGFR-assigned NB-lncRNA markers and reported subtype markers when both luminal breast cancer subtypes were considered together.

We also assessed the overlap between MGFR-assigned NB-lncRNAs markers of each cancer subtype and Seurat-defined markers of normal breast cell subpopulations (Table S18). We found significant overlaps (p-values < 0.001) between markers of basal-like tumors and basal cluster L6, Her2^+^ and both luminal tumors and basal cluster L1 and luminal mature cluster L4 Markers of normal-like tumors were strongly (overlap p-values < 0.005) correlated with all clusters, except L5. Finally, we used the expression levels of 100 MGFR-assigned NB-lncRNA markers (the 20 most frequently assigned to each subtype) to generate a PCA plot of TCGA samples (Fig. 9d). The expression of NB-lncRNA markers successfully distinguished molecular subtypes of breast cancer, with the two first principal components separating them as well as the expression of previously known markers (46) (Fig. 9d). Biomarkers of special interest include the NB-lncRNAs that most contributed with the first principal component to distinguish breast cancer subtypes (Table S18). For example, MGFR-assigned basal cancer subtype marker *DN37008C0G1I10* is a novel isoform of *LINC01956* (located near EN1; Fig. 9c) which is part of a reported triple-negative breast cancer module (70).

## DISCUSSION

Single-cell RNAseq has allowed unprecedented insight into gene expression across different cell populations in normal tissue and disease states. However, almost all studies rely on annotated gene sets to capture gene expression levels, discarding sequencing reads that do not align to known genes. Here, we comprehensively annotated lncRNAs expressed in human mammary epithelial cells, prior to quantitating the transcriptomes of individual cells from healthy breast tissue. On average, we observed ∼900 expressed NB-lncRNAs per cell, compared with ∼5,000 annotated genes. For comparison, when we used the same method to assess the set of confirmed annotated lncRNAs from GENCODE, the average count of expressed genes per cell was ∼1,300. We can attribute this increase in number (compared with NB-lncRNA) to annotated lncRNAs having higher and/or more widespread expression, making them easier to be experimentally detected. Nevertheless, the number of detected lncRNAs is much lower than annotated genes in general, which may reflect the more compartmentalized nature of noncoding transcripts.

The expression of NB-lncRNAs alone could discriminate between luminal and basal cell populations and define subpopulations of both cell types. In the luminal compartment, we observed two luminal progenitor and one luminal mature subpopulation (L0, L2, L4), similar to clustering based on the expression of annotated genes (A1, A2, A4). Two A-clusters (A5 and A7) were defined by a high prevalence of dead cells, based on the presence of mitochondrial markers. As the NB-lncRNAs gene set does not contain mitochondrial genes, the absence of dead cells was expected in L-clusters. Indeed, cluster A5 was virtually absent from the clustering based on NB-lncRNA expression. A subset of cells from cluster A7 was present in basal cluster L7. Despite the absence of Seurat-defined breast epithelial markers, cluster A7 had basal characteristics (e.g. high expression of *CAV1*, *ITGB1* and *TAGLN*), therefore, the resulting L-cluster may have incorporated these cells based on their basal gene expression.

In the basal compartment, while the expression of GENCODE-annotated genes clustered most cells in one heterogeneous cluster of basal cells (A0), NB-lncRNA expression divided the same cells in three subpopulations (L1, L6, L7). This suggests that NB-lncRNA expression may provide an additional layer of information, distinguishing cell subpopulations which annotated genes cannot discriminate. The relevance of this distinction is reflected in the reconstruction of cell hierarchies (Fig. 7d), where myoepithelial cluster L1 is derived from stem-like cluster L3 and a separate lineage emerges from cluster L3 including the intermediate progenitor cluster L5 and basal clusters L6 and L7. When we enforced overclustering of A-clusters, A0 subdivided into two cell pools, one predominantly found in cluster L1 and a second predominantly found in clusters L6 and L7. Notably, cells that form L6 could not be separated from the cells that form L7 based on annotated gene expression alone, even at extremely high resolution. It is important to note, however, that these clusters do not necessarily reflect different cell types but may represent differential cell states or subpopulations of the same cell type that are responding to environmental stimuli. Another difference between A-clusters and L-clusters is that, in the latter, two subpopulations with high prevalence of breast stem-cell markers (A3 and A6) were combined into one cluster (L3), which appeared to be the most stem-like population.

Despite the ability of NB-lncRNAs to cluster subpopulations of mammary epithelial cells, they only poorly distinguish brain cell types, suggesting that the set of lncRNAs expressed in the breast are substantially different from those in the brain. This highlights the need to comprehensively annotate lncRNAs across different tissues and cell types, in a similar way to the annotation of enhancers, which is being performed in a cell type-specific manner. This will be particularly relevant for minor cell populations, which are underrepresented in complex tissues. We suggest that comprehensive transcriptome reconstruction should precede scRNAseq analyses and tissue-specific lncRNAs should be incorporated into the gene set used for clustering. In support of this, in this study alone, we identified >13,600 lncRNAs not found in existing databases, representing nearly 75% of the recovered long noncoding transcriptome of the normal human breast. Several NB-lncRNAs were identified as markers of specific clusters and annotated lncRNAs were also amongst Seurat-assigned markers when only annotated genes were considered (File S4). Lab-based experiments will be required to establish the role of lncRNAs in defining cell types and establishing cell states.

Previous studies have shown that the expression of protein-coding genes in the different breast cancer subtypes reflects their cell-of-origin. For NB-lncRNAs, we confirmed their expression profiles in normal cell subpopulations to be significantly correlated with breast cancer subtypes in TCGA. Moreover, we found Seurat-assigned markers of L-clusters to significantly overlap with the in-house-derived breast cancer NB-lncRNA markers. NB-lncRNA markers of basal-like breast tumors were enriched in markers of basal cluster L6. This may be a reflection of underlying characteristics of basal-like tumors which we observed in cells that form cluster L6, for example higher rates of EMT and a claudin-low profile, which are known hallmarks of basal-like tumors (59,61). Interestingly, all basal L-clusters had gene expression profiles reminiscent of claudin-low type tumors, according to the Genefu classification tool. NB-lncRNA markers of both luminal breast tumor subtypes were enriched in markers of luminal mature cluster L4, which is consistent with previous observations (71), but were also enriched in markers of the basal cluster L1.

To test the capacity of NB-lncRNAs as markers of breast cancer subtypes, we assessed their expression in TCGA samples and derived gene signatures of the different subtypes. We confirmed the NB-lncRNA cancer markers correlated with previously reported protein-coding markers of each subtype and showed their ability to discriminate between breast cancer subtypes. Several lncRNAs previously implicated in breast cancer (Lnc2Cancer (53), lncRNAfunc (54) and the Diermeier *et al* study (55)) were also assigned as markers in our study, including oncogenes *MEG3* and *LINC01089* and mammary tumor associated RNA 18 (*MaTAR18*/*TTC39A-AS1*). *TTC39A-AS1* was also previously found to be a marker of normal luminal mature subpopulations (56) and we identified it as a marker of luminal A and B cancer subtypes. Attesting to the relevance of NB-lncRNAs for future research, ASO-mediated knockdown of *TTC39A-AS1* led to impaired branch development in mammary organoids (55).

It is important to acknowledge the limitations of the present study. For instance, the high prevalence of repetitive elements in lncRNAs may result in inaccurate or incomplete transcript assembly. Individual validation of transcript structures will be required prior to functional studies. In addition, although NB-lncRNA discovery was performed based on bulk sequencing of total RNA, the single-cell RNAseq was performed on oligo-dT primed libraries. This has likely excluded a subset of NB-lncRNAs, which could not be detected at single-cell level. Moreover, the NB-lncRNA discovery was performed using basal and luminal cell populations but other cell types are present in the breast, such as stromal, fat and immune cells. Inclusion of lncRNAs from other cell types will be required to capture the full repertoire of lncRNAs and cell types in normal breast tissue. Adding to the complexity of our study, as cell subpopulations are defined by the expression of protein-coding genes, the labeling of NB-lncRNA based clusters was not straightforward. Finally, as lncRNAs can have lower expression levels, we prioritized the use of data generated by the Fluidigm C1 platform, which results in a higher sequencing depth to the expense of cell number. As RNA sequencing becomes more cost-effective, it will allow the use of 10X Genomics platforms to profile tens of thousands to millions of cells in high depth, expanding our knowledge on lncRNA expression at single-cell level.

In summary, combining tissue-specific lncRNA discovery and single-cell transcriptomics has unraveled new markers of mammary epithelial cell populations and breast cancer subtypes. The fact that a subset of NB-lncRNAs is present in the human breast in both healthy and disease states may further support biomarker discovery and help detect tumorigenesis at the earliest stages. Importantly, although most of our study was performed using Fluidigm scRNAseq data, we observed similar clustering using scRNAseq data generated with the 10x Genomics technology, despite the lower sequencing depth. As 10x Genomics has the fastest growing platform for scRNAseq, this represents an untapped resource for transcriptomics, especially for the study of lncRNAs. Incorporating this novel information could lead to novel diagnostic tools and targets for treatment in a range of cancer types and other complex diseases.

### Data Availability

Raw sequence files were deposited at the SRA under the identifier SUB11631006.

### Code Availability

All the computational scripts used in this work (Script Files 1-12) are available through GitHub at the Functional Genetics Laboratory page, under the repository named ‘NB-lncRNAs’: https://github.com/FunctionalGeneticsLab/NB-lncRNAs. Scripts for bulk RNAseq analysis used for tissue-specificity analyses are available at https://github.com/FunctionalGeneticsLab/GRADE.

## Acknowledgements

The authors thank the donors and their families for the tissues donated for research through the Brisbane Breast Bank and The Wesley Tissue Bank and Dr. William Cockburn for providing reduction mammoplasty tissue. In addition, we would like to acknowledge the QIMR Berghofer HPC team for their support with maintaining the institutional computational infrastructure. We thank Professor Jane Visvader for valuable discussions and critical feedback. We acknowledge the work of Alex Hannah, Fei Wu, Senn Oon and Zach Michaelis in creating a web portal to make NB-lncRNAs available and accessible to the scientific community.

## Author Contributions

JDF and SLE conceived the project and MB designed and directed the study. MB performed the majority of the analyses. IPA and ISR critically reviewed the computational code and manuscript, IPA created and maintains the GitHub repository, ISR performed the RAMPAGE and splice site analyses and deposited sequencing data to public repositories. SL provided the normal breast samples, which were pre-processed by KF. WS processed the normal breast samples from the Brisbane Breast Bank. SLE made the bulk RNA sequencing libraries. MB, JDF and SLE interpreted the data and co-wrote the manuscript with input from all authors.

## Supplementary information

Supplementary files, tables and figures are supplied separately.

