## Supplementary Material for "Redefining normal breast cell populations using long noncoding RNAs"

**Running Title: Noncoding RNAs are markers of breast epithelial cell types.**


**Authors**

Mainá Bitar[1], Isela Sarahi Rivera[1,2], Isabela Pimentel de Almeida[1], Wei Shi[1], Kaltin Ferguson[3], Jonathan Beesley[1], Sunil R Lakhani[3,4], Stacey L Edwards[1*], Juliet D French[1*]

**Affiliations**

[1] Cancer Program, QIMR Berghofer Medical Research Institute, Brisbane, 4029, Australia.

[2] School of Biomedical Science and Institute of Health and Biomedical Innovation, Faculty of Health, Queensland University of Technology, Brisbane, 4001, Australia.

[3] UQ Centre for Clinical Research, Faculty of Medicine, The University of Queensland, Brisbane, 4029, Australia.

[4] Pathology Queensland, The Royal Brisbane & Women's Hospital, Brisbane, 4006, Australia.

* These two authors contributed equally to this work

**Correspondence:**

+ 61 7 3362 0222

**SUMMARY OF SUPPLEMENTARY INFORMATION**

**1) Description of Supplementary Files:**

**File S1:** *Sample preparation, sequencing and transcriptome assembly.*
Contains detailed methods and results for every step of the de novo assembly pipeline used for transcriptome reconstruction and subsequent analysis to identify lncRNAs and confirm transcript structure using RAMPAGE and long-read sequencing data.

**File S2:** *Internal databases and retrieved public data.*
Contains information on the source and composition of in-house databases for known lncRNAs and normal breast cell markers. Also contains details on the downloaded brain single-cell RNA sequencing (scRNAseq) data.

**File S3:** *Functional characterization of NB-lncRNAs.*
Contains further information regarding the characterization of regulatory NB-lncRNAs, such as enhancer-derived, promoter-associated and terminus-associated lncRNAs.

**File S4:** *Single-cell RNA sequencing analysis.*
Contains detailed methods and results for the scRNAseq analyses. While the main findings are presented in the manuscript, additional analyses and results are presented in the file, along with further details on the clustering experiments. Also contains a list of NB-lncRNA markers of interest and NB-lncRNAs associated with housekeeping genes.

**File S5:** *Replacement of SCENT gene network for potency assignment*
Contains the methods to construct a coding <-> noncoding gene network which we used to replace the default SCENT gene network. The in-house network was built using LncRNAs2Pathways data as described.

**2) Description of Supplementary Tables:**

**Tables S1-S18:** Each supplementary table includes a tab named “Table Description”, in which all of its tabs are described. Supplementary tables are provided separately as individual files.

**3) Description of Supplementary Figures:**

**Figures S1-S6:** Supplementary figures are provided at the end of this file, with their corresponding legends.

**SUPPLEMENTARY: SAMPLE PREPARATION, SEQUENCING AND TRANSCRIPTOME ASSEMBLY**

**A) *Preparation of normal breast samples***

Normal breast tissue samples were collected during reduction mammoplasties of five healthy donors [1] (Table S1) and processed as per previously described [2]. In brief, samples were digested into single-cell suspensions using collagenase and flow cytometric sorting was performed with BD FACSAria III to isolate distinct subpopulations. To enrich for epithelial populations, suspensions were stained with a lineage marker antibody combination of anti-CD31, anti-CD45 and anti-CD140b, designed to exclude endothelial, hematopoietic, and leukocyte cells (respectively; Table S1). The remaining lineage-negative cell superpopulation was sorted into basal (EpCAM$^{low}$/CD49f$^{hi}$), luminal progenitor (EpCAM$^{hi}$/CD49f$^{hi}$), and luminal (EpCAM$^{hi}$/CD49f$^{low}$) cell types. Total RNA was extracted from pooled cells, with each replicate yielding 1.0μg to 3.0μg of high-quality RNA (RIN ≥ 9.0, measured in Agilent 4200 Tapestation).

**B) *Sequencing of normal breast samples***

Samples were sent to NovogeneAIT Genomics (Singapore) for ribosomal RNA depletion (Ribo-Zero™ Plus rRNA depletion kit), total RNAseq library construction (NEBNext® Ultra™ Directional RNA Library Prep Kit) and sequencing using Illumina Novoseq 6000. Fragment size was 280 and reads were sequenced as pairs of 150nt each read. Each sample was sequenced across 2-3 lanes, yielding ~30-67 million read pairs (average 49.9 million). Notably, others have shown that *de novo* transcriptome assembly performed by Trinity on 50 million reads could unequivocally assign transcripts even to genes that have very low expression levels, deeming our acquired sequencing depth ideal for gene discovery [3]. The percentage of duplicate reads per sample was on average <60% (which is expected for the high depth of sequencing, e.g. Galaxy Project; website reference[1]). Average Phred-scale quality for each sample was between 37 and 39, which represents a base call accuracy of approximately between 99.95% and 99.99%. Quality metrics were calculated using FastQC v0.11.5 and are shown in Table S2.

**C) *Pipeline structure and detailed results of transcript reconstruction***

**1. Pre-processing of reads to remove contaminants**

*Overview*

The first part of the in-house pipeline was designed to optimize the sequencing reads that serve as material for the *de novo* assembly. This includes performing quality control routines before and after read processing, correcting and trimming reads, removing ribosomal reads and assessing the resulting 'clean' reads to confirm strandness and rule out contamination. This part of the pipeline was largely based on Harvard's 'Best Practices for *de novo* Transcriptome Assembly' guidelines (Harvard Informatics; website reference[2]).

*1.1 Error correction*

*Methods:* Sequencing errors in RNAseq reads complicate *de novo* assembly by creating artificial nodes in De Bruijn graphs during isoform resolution [4]. Rcorrector [5] is a top performing method to correct random sequencing errors in Illumina RNAseq reads. It places k-mers derived from input reads in a *trust scale*, based on prevalence. Low-frequency k-mers are converted to their high-frequency counterpart. FUR-PE (Adam Freeman; website reference[3]) is an accessory tool used to further remove all reads deemed *uncorrectable* (typically low complexity sequences), fix the format of the header for compatibility with downstream tools and report a summary of read correction. We used Rcorrector v.1 followed by FUR-PE v.2016. *Results:* Average read quality measured in the Phred scale increased from 38.6 (raw reads; Table S2) to 40.3 (corrected and trimmed reads; Table S3), representing a decrease in the calculated error rate from 1 base in ~7,000 to 1 in ~10,000. Between 39.8% and 59.9% of the reads in each sample were corrected using Rcorrector (average of 52%; Table S4). *Uncorrectable* reads were filtered using FUR-PE, with 5.8-16.8% of the read pairs being removed per sample (average of 9.5%; Table S4). Nevertheless, to ensure our choice of correcting the reads before assembly was the best decision, we kept a separate copy of uncorrected reads and compared results at the end of the pipeline.

*1.2 Trimming*

*Methods:* Corrected reads were trimmed with Trimmomatic v.036 [6] to remove Illumina adaptors. To balance read length and allowed errors, we set the *MAXINFO* flag to 100:0.4, maximising the value of each read. A low value of this flag (<0.2) would favour longer reads and a high value (>0.8) would favour read correctness.

*Results:* Between 67% and 79% of the read pairs survived trimming (average of 72%, Table S4) and an additional 18% to 30% of the forward reads survived independently (only 0.5% to 1.6% of the reverse reads survived independently). Uncorrected reads were also trimmed, for control, with between 60% and 71% of the pairs surviving trimming (average of 64%, Table S4) and an additional 25% to 37% of the forward reads surviving independently (0.5% to 1.5% of the reverse reads surviving independently). The average Phred score across reads was raised to 40 or higher in all samples, as a consequence of read treatment.

*1.3 Ribosomal RNA filtering*

*Methods:* We used the Ribo-Zero™ Plus rRNA depletion kit to deplete rRNA in the samples during library preparation. Reads were also computationally filtered to remove remaining rRNA-derived reads using BBduk based on a set of 169 human rRNA sequences (Table S8, mainly derived from the Silva database [7].

*Results:* A high proportion of rRNA-derived reads is expected in RNAseq experiments [8]. On average, samples contained 35% rRNA-derived reads pre-filtering (median was 26% and range was 13%-67%, Table S4). For comparison, the best possible depletion method (using rRNA-excluding random hexamers during sequencing) can only lower the fraction of rRNA in samples to 13% [9].

*1.4 Additional quality tests*

Using the *infer_experiment* routine of RSeQC v.2.6.4 [10], we confirmed that 93%-96% (average of 94%) of the sequenced reads were coming from one strand only (based on fraction of reads explained by "1+-,1-+,2++,2--", which stands for "FR as first strand"). Reads were aligned to the GENCODE (GCRh38 release 79) version of the reference human genome using Bowtie2 v.2.2.9 [11] to rule out external contamination. Overall alignment rates varied from 80% to 88% per sample (average of 84%), as expected for high-quality RNAseq experiments with human samples [12]. Using the *multiBamSummary*, *plotCorrelation* and *plotPCA* functions of DeepTools v.3.5.0 [13], we generated scatter plots of pairwise Pearson correlations and a clustered heatmap of all samples' correlations, which revealed an overall absence of batch effects (samples clustered by cell type rather than donor). All sample correlations (measured by Pearson coefficients) were higher than 0.7, with an average of 0.9.

### 2. *De novo* assembly of transcripts

*Overview*

At the second part of the designed pipeline, we used Trinity v.2.8.4 [3] to perform the *de novo* assembly of transcripts and additional methods to ensure its quality. Transcripts were further filtered to generate a refined version of the assembly. Trinity is a top performing and the most widely used method for *de novo* transcriptome assembly (e.g. [14], PMID: 27054874, PMID: 25759274). BUSCO v.20161119 [15] and TransRate v.1.0.3 [16] are the UC Davis recommended tools for assembly assessment (UC Davis Data Intensive Biology Program; website reference[4]), which we used after Trinity to assess completeness and accuracy. BUSCO (Benchmarking Universal Single-Copy Orthologs) is a method to measure transcriptome completeness, based on an internal database of known near-universal single-copy orthologs. By confirming the correct assembly of these genes, one can expect the rest of the transcriptome to be equally well resolved. TransRate is a tool to score assemblies for their quality, based on experimental evidence (e.g. the sequenced reads). In addition, TransRate can filter lower quality transcripts.

*2.1 Trinity* de novo *assembly*

*Methods:* Briefly, Trinity works by generating contiguous overlapping sequence segments (*contigs*) based on k-mer overlaps. Each final *contig* represents a transcript isoform of a given gene and the set of all generated *contigs* is a compilation of the transcriptome being studied.
*Results:* We combined all available treated reads (from 'part 1') in two unique Fasta files (Trinity *left* and *right* read files) as input for the *de novo* assembly. Over 270,000,000 reads were submitted, 93% of which used by Trinity to generate transcripts. The initial assembly comprised 868,155 genes and 1,047,401 transcripts. It had an N50 of 1,065 and average transcript length of 683 with GC% of ~48 (Table S5), which is intermediate between the reported human GC% of coding (~52%) and noncoding (~44%) isoforms [17].

*2.2 Quality assessment of the raw assembly*

*Methods:* We used BUSCO based on Augustus 3.2.2 and odb9 and both eukaryotic and mammalian ortholog sets to assess assembly completeness. We used TransRate to assign scores to the assembly, based on the proportion of supporting input reads and other metrics.
*Results:* More than 99% of the eukaryotic (301/303) and 86% of the mammalian (3522/4104) conserved orthologs were recapitulated in the assembly (Table S5). According to TransRate, the assembly had an optimized score of 0.36 and overall score of 0.18. It reaches a 65% good-mapping rate and 32% human genome coverage. A *de novo* assembly with this optimized is better than 50% of published *de novo* assembled transcriptomes that have been deposited in the NCBI TSA0 [16], which is considered a very high quality overall.

*2.3 TransRate filtering*

*Methods:* TransRate was also used to filter out contigs that fail to meet the minimum individually quality suggested by the method.
*Results:* In total 627,743 transcripts and 535,696 gene elements (>60%) passed the quality cutoff, increasing the N50 to 1,383 and the average contig length to 826 (Table S5).

*2.4 Overall assembly score*

*Methods:* Recently, Hölzer and Marz [14], assessed 10 of the best and most used *de novo* assembly tools using 20 selected quality metrics. Trinity was ranked second, with an overall score of 12.4 versus 14.2 obtained by the top performing tool. Authors used data from human samples, sequenced as 101nt paired-end reads with a strand-specific protocol, which is comparable to the strategy we used. Of note, while our samples reached an average depth of ~50 million reads, samples in their study reached 100 million reads per sample. We assessed 9 of the 20 metrics and compared our assembly with their published results.

*Results:* Our optimized in-house assembly performed better than the reported Trinity assembly for 7 of the 9 metrics (Table S5), the exceptions being the percentage of transcripts with identified open reading frame (ORF) and the N50 (while our in-house assembly had 42% ORF percentage and N50 1,383 the published Trinity assembly had respectively 51% and 1,840). Our in-house assembly performed better than the top-ranked tool in the study for reference coverage and two BUSCO metrics, better than the second-ranked for number of transcripts longer than 1kb and TransRate optimal score and better than the third-ranked tool for overall mapping rate and percentage of uncovered bases in the human transcriptome. We recalculated the overall score of the top performing tool regarding these 9 metrics (substituting the Ex90N50 for the N50) as 5.95 while our in-house assembly reached a score of 6.66, thus surpassing the best tool by >10% (Table S5).

*2.5 Splicing assessment*

*Methods:* Gmap v2020-09-12 [18] was used to map assembled transcripts to the GENCODE reference human genome. Aligned transcripts were further characterised as spliced or unspliced according to the presence of 'N' tags. After consulting the literature, we defined a minimum intron size of 50bp to represent human transcripts [19,20].

*Results:* Approximately the same fraction of transcripts (~400,000 and ~250,000, or ~40% each) in the raw and optimized assemblies map to 1-2 locations in the human genome. Nearly half of the remaining align to three genomic locations, while the rest aligned to more. In the optimized assembly, ~90,000 transcripts were classified as spliced and ~537,000 as unspliced.

*2.6 Redundancy reduction and read support requirements*

*Methods:* We used the *align_and_estimate_abundance.pl* script from Trinity to assess read support in terms of normalised number of reads per transcript (FPKM). Two filters were applied, a more permissive one, of 0.5 FPKM for multiexonic and 3 FPKM for monoexonic transcripts, and a stricter one, of respectively 1 FPKM and 5 FPKM. We assessed the presence of redundant transcripts using CDhit v.4.6.8 [21] with either 95% or 98% identity cutoffs and >90% coverage cutoff and found less than 200 transcripts to be redundant, which we deemed negligible.

*Results:* From the ~90,000 spliced and ~537,000 unspliced transcripts, respectively ~85,000 (92%) and ~300,000 (56%) passed the more permissive FPKM cutoff for read support. Of note, while the number of supported spliced transcripts was approximately the same applying the stricter FPKM cutoff (~80,000, i.e. <5% decrease), the number of unspliced transcripts was nearly halved (~160,000, i.e. ~90% decrease). The ratio of monoexonic to multiexonic transcripts in the assembly is therefore of between 3 and 2 to 1. The 384,182 transcripts comprise the final filtered assembly, or the comprehensive transcriptome of normal breast, in which we performed noncoding transcript discovery.

### 3. Identification of noncoding isoforms and lncRNAs

*Overview*

The last step of the pipeline identifies the lncRNA from the pool of assembled transcripts. We used ezLNCpred v.1.0 [22] for coding potential calculations and, in parallel, FEELnc v.0.2 [23] to identify lncRNAs, overlapping both transcript sets. The Python package ezLncPred is a comprehensive suite for noncoding transcript identification, integrating multiple state-of-the-art prediction models. FEELnc is an implementation of a machine learning algorithm capable of identifying lncRNAs based on sets of high-confidence protein-coding and lncRNA transcripts of the same species. Others have reported lncRNA predictions obtained by combining FEELnc and coding potential calculators to yield robust results [24].

*3.1 Prediction of coding potential*

*Methods:* Four programs were used to predict the coding potential of all transcripts: CPAT, CNCI, CPC2 and PLEK. All were independently run using the ezLNCpred environment.

*Results:* Coding potential was detected for 83,034 transcripts by at least one tool. The noncoding counterpart has 374,984 transcripts and the overlap is due to inconsistency between tools (Table S6). A total of 198,926 transcripts (52%) are classified as either coding or noncoding by different tools, while 182,827 (46%) are unambiguously classified and 1% fail to be classified. From the unambiguously classified, ~97% (176,784 of 182,827) are noncoding transcripts detected by all four tools. On average, all four tools agree over nearly half of all predictions (176,784 of 374,984, or >47%) of noncoding potential and less than 10% of the coding predictions (6,043 of 83,034 or ~8%). This indicates that methods seem to agree more on ORF absence than presence. Less than 7% of all transcripts with no detected coding potential were individually predicted by only one tool. In total, 176,056 transcripts were predicted by two or more tools as noncoding and not by any tool as coding. We decided to consider all 346,324 transcripts predicted as noncoding by at least two the tools (Table S6).

*3.2 lncRNA identification*

*Methods:* We sourced protein-coding and lncRNA transcripts from the set of GENCODE-annotated genes. For higher confidence, we eliminated entries labelled as 'TEC' (i.e. to be experimentally confirmed). Therefore, all the supplied transcripts are assumed to have been experimentally validated by the consortium. FEELnc is divided in three modules that sequentially discard non-lncRNA transcripts ($FEELnc_{filter}$), identify lncRNAs from the remaining subset ($FEELnc_{codpot}$) and classify the lncRNAs based on their genomic context and nearest annotated genes ($FEELnc_{classifier}$). The second module is the most complex and includes the Random Forest algorithm that classifies the training sets of reliable protein-coding and lncRNAs provided by the user to adapt the prediction scores for each analysis.

*Results:* FEELnc predicted 30,722 transcripts to be lncRNAs (Table S6), with 90% supported by two or more of the ORF detection strategy presented in '3.1'. Notably, only 900 transcripts were not supported by CPAT, the method that most agreed with FEELnc predictions. To minimize false positives, we filtered out the remaining 10% unsupported transcripts, leaving 27,553 high-confidence lncRNA candidates (which we then named normal breast lncRNAs or NB-lncRNAs). From these, 17,390 (~65%) are spliced and 10,163 (~35%) unspliced. Moreover, 12,704 are sense exonic, 10,468 sense intronic, 4,428 intergenic and 3,122 antisense gene-overlapping (Table S6). Nearly 80% of these transcripts (21,496) have additionally passed the stricter FPKM cutoff for read support (1 and 5 FPKM, see '2.6').

*3.3 Previously annotated lncRNAs*
*Methods:* We aligned the obtained lncRNAs to the human transcriptome (the hg38 version of the GENCODE annotation) and to the in-house database of known lncRNAs (Table S8). We defined as 'previously annotated' the lncRNAs candidates with at least 90% sequence identity and 75% reciprocal coverage to known transcripts.
*Results:* Only <15% (3,642 of the >27,500) of the assembled lncRNA candidates coincided with annotated transcripts (Table S9). From these, 80% are mapped to transcripts that are not annotated as protein-coding. Considering the annotated transcript that best represented the assembled lncRNA in terms of sequence coverage and identity, 598 were intron-derived transcripts, 590 antisense transcripts, 454 intergenic lncRNAs and 413 transcripts nonsense mediated decay targets. In addition, 1,116 assembled lncRNAs coincide with ncRNAs from the in-house database, 247 of which lack GENCODE annotation (Table S9). In total, 3,889 assembled lncRNA candidates were known previously, 751 being possible false-positives (as the related annotated genes are exclusively protein-coding). Overall, ~90% of the assembled lncRNAs are either novel transcripts or related to previously annotated noncoding transcripts. Splicing was detected in ~95% of all the annotated lncRNA candidates and in ~50% of the unannotated counterpart. This discrepancy seems to demonstrate an overrepresentation of spliced transcripts in the current human transcriptome, which may be a consequence of the common practice of dismissing monoexonic transcripts regardless of read support.

**D) *Inclusion of monoexonic transcripts***
Liu and collaborators [25] found >75% of lncRNAs to be single-exon when performing *de novo* assembly of transcripts from the human brain. Assessing entries in the in-house database of known lncRNAs, we found nearly half the transcripts to be spliced and the other half unspliced (respectively 51% and 49%). For example, over 60% of the genes in the MiTranscriptome are unspliced. Moreover, more than 70% of the intergenic (5,968/8,043) and antisense (4,197/5,816) lncRNAs in the reference genome have only one annotated transcript, indicating lack of complex splicing. For sense-intronic genes, the proportion surpassed 95%. Protein-coding genes have the opposite behavior, with just 25% (5,465/21,855) of the annotated genes lacking evidence of splicing. This inversion on the proportion of spliced to unspliced transcripts in protein-coding versus noncoding genes supported our decision not to filter out monoexonic elements. In agreement, Diermeier and collaborators [26] have previously assessed lncRNAs in mammary glands and breast cancer and found only 10% of these to have complex splicing structures with multiple exons.

**E) *Confirmation of the transcript structure of NB-lncRNAs***

***Overview***
To test the performance of our transcript reconstruction strategy, we assessed the full-length read support and transcription start site (TSS) of the assembled NB-lncRNAs.

**1. Confirming the TSSs of NB-lncRNAs**

*Methods:* Normal breast epithelium RAMPAGE data was obtained from the ENCODE portal (ENCSR909QWB and ENCSR598TAK from the Thomas Gingeras Laboratory; Website Reference[5]). Samples are from two female individuals, ages 51 and 53, both of which died from cerebral vascular accidents. Mammary tissue was collected less than 8 hours after death in both cases and sample descriptions were as follows: (i) 50-70% fat content, remainder is stroma with benign ducts and lobules and (ii) 10% fat content, lobular and ductal elements in predominantly fibrous stroma. Authors declare that RAMPAGE libraries were built from rRNA-depleted total RNA >200 nucleotides in size and sequenced as stranded paired-end Illumina Hi-Seq reads of 101nt. We merged all samples, obtaining 21,998 TSS peaks. Additional methodological details can be obtained from the ENCODE portal. We also used in-house RAMPAGE data of breast cancer cell lines BT549, MCF10A, MDAMB231 and SUM149. Briefly, libraries were sequenced as 150bp paired-end reads and demultiplexed using the *icetea* library in R before being aligned to the reference genome (GENCODE GRCh38 v34) supplemented with the NB-lncRNAs using STAR v. 2.7.1a [12]. RAMPAGE peaks were called using the *call_peaks* script in GRIT v2.0.4 (website reference[6]). We used Bedtools v.2.29.0 to intersect TSS peaks with NB-lncRNA coordinates, and requested that the TSS should be located upstream the lncRNA transcript and no more than 500bp away. Additionally, when the TSS peak was internal to the predicted lncRNA, only overlaps of up to 50bp were accepted.
*Results:* We confirmed the TSS of 960 (approximately 5%) of our NB-lncRNAs in high resolution (<500bp upstream or <50bp internal to the predicted first base) using public ENCODE data. For comparison, 10,259 (approximately 15%) annotated genes had their TSS confirmed with the same dataset. Using the in-house RAMPAGE datasets, we confirmed 1,443 additional TSSs, 319 from MCF10A data, 603 from SUM149, 860 from BT549 and 931 from MDAMB231. In total, considering both publicly available and in-house generated TSS annotations, we confirmed 1,810 NB-lncRNA start sites (Table S7).

**2. Full-length read support to NB-lncRNAs**

*Methods:* We downloaded 12 long-read files from the NCBI's SRA (PRJEB44348 and PRJNA522784; website reference[7]), 11 from MCF7 breast cancer cells [27] and one from MCF10A cells [28]. Samples were sequenced in the Oxford Nanopore Technologies' MinION platform, resulting in 30 thousand to 10 million long-reads (2.7 million average) and average read lengths from 180nt to nearly 1,400nt. Additionally, we generated 33.1 million long-reads from SUM149 breast cancer cells at an average length of 1,184 nucleotides. Total RNA was sent for long-read cDNA sequencing by PromethION to the Garvan Institute Nanopore Sequencing Facility (Australia). We used BLAT v.35 [29] to report all alignments between assembled transcripts and each sequence in the long-read libraries with at least 80% sequence identity (set low, to accommodate the high error rate which is common to long-reads). BLAT was allowed to run for 300 CPU hours, which exhausted all alignment possibilities for NB-lncRNAs in the MCF7 and MCF10A libraries but not for the SUM149 library, which was sequenced to a greater depth. We then used an in-house Bash script to recover high confidence aligned pairs. High-confidence pairs were defined by >70% reciprocal coverage and <10% indels in aligned regions.
*Results:* With >30,000 high confidence pairs recovered, 940 lncRNAs supported by long-reads of MCF7 or MCF10A cells and 752 by long-reads of SUM149 cells. In total we confirmed 1,310 NB-lncRNAs by long-read support (Table S7).

### ***Website References:***


1: https://training.galaxyproject.org

2: https://informatics.fas.harvard.edu/best-practices-for-de-novo-transcriptome-assembly-with-trinity.html

3: https://github.com/harvardinformatics/TranscriptomeAssemblyTools/blob/master/FilterUncorrectabledPEfastq.py

4: http://ivory.idyll.org/lab

5: https://www.encodeproject.org/rampage

6: https://github.com/nboley/grit

7: https://www.ncbi.nlm.nih.gov/sra

**SUPPLEMENTARY: INTERNAL DATABASES AND RETRIEVED PUBLIC DATA**

**A) *In-house dataset of known lncRNAs***

We generated an in-house dataset comprising >112,000 entries from the following sources: BIGtranscriptome (part of lncRNAKB [30]; Cabili *et al*. 2011 [31]; CancerSEA [32]; Lanzos *et al*. 2017[33]; LNCipedia (part of lncRNAKB); LncRNADisease [34]; MiTranscriptome (part of lncRNAKB) and RNAcentral [35]. Data from BIGtranscriptome, LNCipedia and MiTranscriptome have been previously integrated and made available as the lncRNA knowledgebase (lncRNAKB), which contains over 77,000 human lncRNAs. The LNCipedia database [36] itself combines data from the lncRNAdb (105 literature-retrieved transcripts), the Broad Institute human body map (14,279 lincRNA transcripts), ENSEMBL (version 99 at the time of our download, comprising 25,075 transcripts), NONCODE (93,164 transcripts), GENCODE (version 13 with 19,812 transcripts), RefSeq *biomol_ncrna_lncrna* entries (2014 version with 4,774 transcripts and release 106 with 5,487 transcripts), Nielsen *et al*. transcriptome data from 12 human tissues (7,656 transcripts), Hangauer *et al*. data from the *de novo* assembly of ~7 million transcripts (5,339 curated lncRNAs with FPKM>1), Sun and Gadad *et al*. GROseq and RNAseq analysis of MCF7 transcriptomes (2,305 transcripts), FANTOM CAT lncRNAs (27,719 minus 34 transcripts that were in conflict with the HUGO gene boundaries). The BIGtranscriptome contains 105,494 known and 8,692 novel genes, ncRNAs from Lanzos *et al*. add to 5,914 genes, being 45 cancer-related and the MiTranscriptome contains 175,772 lncRNA transcripts.

We filtered the lncRNAKB database to exclude noncoding genes that are not lncRNAs (keeping 73,611 genes) and genes from chromosomes that are not fully assembled (keeping 72,166 genes). We then used Bedtools v.2.26.0 to merge genes on the same strand which were adjacent or contained within one another, resulting in a final dataset of 51,524 genes of between 3 and 1,375,316 nucleotides in length. We then combined entries from all the mentioned resources, filtering by length (between 200 and 25,000 nucleotides) and 'lifted' the genomic coordinates to the hg38 version of the human genome (when needed), using the UCSC browser portal (website reference[1]). The resulting list includes 2,528 annotated lncRNAs, 2,620 lncRNAs from Cabili *et al*., 31,389 lncRNAKB entries and 184,182 RNAcentral entries. After reducing redundancy, the final set contained 112,438 lncRNAs (Table S8).

**B) *Database of normal breast epithelial cell markers reported in the literature***

We thoroughly reviewed the literature followed by information retrieval to find markers of normal breast cell populations, preferably derived from single-cell RNA sequencing (scRNAseq) studies. We selected 11 publications (Table S8), all except one listing at least ten genes. Additionally, we retrieved markers of 11 breast cell populations available in the CellMarker database [37] to add 67 genes to our in-house dataset (another 24 genes were already listed in the selected publications). In total, 79 genes in the dataset are markers of stem/precursor cell populations, 58 are markers of basal/myoepithelial cell populations (32 exclusively), 110 of luminal cell populations (64 exclusively) and 28 encode proteins involved in milk production. The final database comprises 366 genes, which we divided into 26 classes of at least 3 genes each (Table S8).

**C) *Single-cell RNA sequencing data for brain cells***

The transcriptomes of 466 brain cells were obtained from the SRA database (PRJNA281204 [38]). Cells were distributed as below, according to authors' labels provided.

| Cell Type | Number of Cells |
|---|---|
| Astrocytes | 62 |
| Endothelial | 20 |
| Fetal quiescent | 110 |
| Fetal replicating | 25 |
| Hybrid | 46 |
| Microglia | 16 |
| Neurons | 131 |
| Oligodendrocytes | 38 |
| OPC | 18 |
| *Total* | *466* |

Markers for each cell type are also listed by authors (below) and were used to characterize the different cell clusters obtained based on GENCODE-annotated gene expression. (1) Astrocytes: *SLC14A1, GLIS3, GLI3, PPP1R3C, CHRDL1, CYBRD1, CTH, SORCS2, ITGB4, RNF43, NWD1, PAQR6, C16orf89, ALDH1L1, TRIM66, HGF, CBS, ITGA7, SLC30A10, SLC4A4, FGFR3, BMPR1B, ATP13A4* and *AQP4*. (2) Endothelial cells: *APOLD1, TM4SF1, FLT1* and *A2M.* (3) Microglia: *GPR183, CCL4, CD83, LAPTM5, CSF1R, HLA-DRA, BCL2A1, CD14* and *CCL2.* (4) Neurons: *KCNK1, KIAA1324, LNX1, NELL1, COBL, SLITRK1, DPYSL5, C14orf37, DLX1, DLX2, DLX5, DLX6, GLRA2, SLC10A4, EGFR, SST, PNOC, NXPH1, BCL11A, DCN, TMEM130, CNTN4, CDO1, NFASC, LRRTM3, GRIA3* and *RELN.* (5) Oligodendrocytes: *DAAM2, ASPA, MAL, SEC14L5, MAP6D1, DPYD, PPP1R14A, GJB1, FA2H, MAG, CDK18, LGI3, SHC4, UGT8, KLK6, KCNH8, GPR37, MOBP, LPAR1, ADAMTS4, ERMIN, OPALIN, CLDN11, PLEKHB1, GSN, GRM3, CNP, MBP* and *PLP1.* (6) OPCs: *PDGFRA, LHFPL3, MEGF11* and *PCDH15.*

**D) *UMAP versus tSNE as graphical representations of single-cell clustering***

There are two main differences between UMAP and tSNE clustering methods: (1) the initialization of each algorithm and (2) their cost function for dimension reduction. While UMAP initializes the algorithm using a graph Laplacian, tSNE uses a random initialization method. Moreover, UMAP uses cross entropy as a cost function for dimension reduction and tSNE implements the Kullback-Leibler (KL) divergence cost function, which cannot preserve global distances without becoming very computationally expensive. Oskolkov (Lun University's Science for Life Laboratory) further discussed the main differences between UMAP and tSNE (website reference[2]). We decided to use UMAP in our analyses, mainly due to its non-randomized initialization and capacity to ensure that points placed distantly in high dimensions will remain distant in low dimensions.

***Website References:***


1: https://genome.ucsc.edu/cgi-bin/hgLiftOver
2: https://towardsdatascience.com/tsne-vs-umap-global-structure-4d8045acba17

### SUPPLEMENTARY: FUNCTIONAL CHARACTERIZATION OF NB-lncRNAs

#### A) *Characterization of regulatory NB-lncRNAs*

##### *Overview*

As many lncRNAs act by regulating the expression of nearby protein-coding genes, we characterized regulatory NB-lncRNAs by identifying transcripts originating from enhancer elements, promoter regions or terminal untranslated regions (UTRs) of annotated genes. Enhancer-derived NB-lncRNAs (NB-elncRNAs) can recruit transcription factors and act in both short-range (*cis*) and long-range (*trans*). Promoter-associated noncoding RNAs (pancRNAs) are transcribed from the vicinity of a TSS and often modulate the protein-coding gene expression [39]. Terminus-associated lncRNAs (TALRs) are transcribed from the UTRs of mRNAs and can also regulate their expression. TALRs typically originate from the 3' end and are often independently transcribed [40].

##### 1. Overlaps between NB-lncRNAs and enhancers

*Methods:* To identify NB-elncRNAs, one million enhancers were obtained from normal epithelial breast cells and breast cancer cell lines. We downloaded data for *cis*-regulatory elements of epithelial breast samples from the ENCODE portal (ENCFF753NXE and ENCFF806DLU; website reference[5]) and cell-specific enhancers of MCF7, MCF10A, MDA-MB-231 and T47D breast cancer cells from the EnhancerAtlas 2.0 [41]. All these subsets together added to 1,049,094 elements, which are largely non-redundant (merging nearby elements allowing a maximum 10bp distance between ends would result in ~850,000 elements). High-confidence NB-elncRNAs were defined as those co-localised with enhancers with at least 70% mutual overlap.

*Results:* We found 349 elncRNAs, transcribed from 352 annotated enhancers, being 97 from ENCODE and 255 from EnhancerAtlas. From the latter, 127 enhancers were from MCF7, 64 from MCF10A, 37 from MDA-MB-231 and 31 from T47D cells (4 were shared by 2-3 cell types). Authors of EnhancerAtlas developed and validated the EAGLE method [42] to predict target genes for enhancers. Fifteen of the 255 enhancers had resolved targets, varying from only one annotated gene target to 69 (average 18). Over half (193 NB-elncRNAs) were consistently expressed at 1 TPM or higher in at least one breast cell type. Four breast epithelial markers (Table S8) had NB-elncRNAs transcribed from an associated enhancer. One example is NB-elncRNA DN55837C0G1I1 (Fig. 1e), which was consistently expressed in both luminal cell subpopulations, and transcribed from an enhancer located in an intron of *EDN1* (a regulator of cell proliferation in cancer).

##### 2. Overlaps between NB-lncRNAs and promoters

*Methods:* EPDnew [43] is an experimentally validated database from the Swiss Institute of Bioinformatics (SIB). The curated list contains 29,598 human promoters, which we intersected with all NB-lncRNAs. We defined promoter regions as the 500 nucleotides upstream of the TSS for each gene and retrieved all lncRNAs overlapping these regions, varying the minimum requested coverage of the promoter region from 10% to 80% (i.e. from 50nt to 400nt).

*Results:* The number of lncRNAs overlapping promoter regions on either sense or antisense directions was 2,730 at a 10% coverage cutoff and 1,968 at 80% coverage cutoff which we analysed in more detail. The NB-pancRNAs transcribed from the promoters of 1,686 protein-coding genes (Table S10). NB-pancRNAs are transcribed from the promoters of 39 known breast markers, including *CAV1, CCL2, ELF3, ITGB1, MYLK, SYTL2, TCF4, VIM* and *ZEB1*. One example is DN61326C0G1I2, which is exclusively expressed in luminal progenitor cells, is a pancRNA of *ACHE*, a gene that is commonly deleted in tumors with *ERBB2* amplifications [44] (Fig. 1f).

#### 3. Overlaps between NB-lncRNAs and UTRs

*Methods:* To find terminus-associated lncRNAs, or TALRs, we downloaded 3'UTRs and 5'UTRs sequences from two different sources: (1) AURA, or the Atlas of UTR Regulatory Activity and (2) the UTRdb [45,46]. Combined, these resources have approximately 500,000 UTRs, although we did not assess redundancy. We used BLAT to align the newly discovered lncRNAs to this set of UTRs and further filtered the outputs to requiring the UTR region to be longer than the TALR and using either 95% minimum or complete coverage of the lncRNA and at least 20% coverage of the UTR.

*Results:* We found from 272 to 825 TALRs, depending on the minimum UTR coverage being requested. Considering only AURA entries, which have the specification of being 3' or 5' UTRs, we estimate nearly 25% of the TALRs to be derived from 5' ends and the remaining >75% to be derived from 3' ends, which is in agreement with the current knowledge on TALRs. The 825 TALRs target 731 annotated genes, from which 717 are protein-coding (Table S10). NB-TALRs were found associated with 11 normal breast marker genes (*CAMK2D, CCND2, CREB5, CREB3L2, CSNK1A1, GSK3B, PDGFRA, PLB1, RGS5, SOX9, VEGFA*). One example is NB-TALR DN124180C1G1I1, which was uniquely expressed in basal cells and spanned the 3'UTR of the basal marker *CCND2* (Fig. 1g).

### B) *Annotated breast cancer-related NB-lncRNAs*

Ninety-eight NB-lncRNAs have been previously implicated in breast cancer by the curated databases lncRNAfunc [47] and Lnc2Cancer [48]. Overall, we identified nearly 25% of the known experimentally-supported breast cancer-related lncRNAs in these resources (48/255 from lncRNAfunc and 63/193 from Lnc2Cancer). The list includes well-known oncogenes such as *CASC5*, *CRNDE*, *DANCR*, *GAS5*, *HOTAIR*, *LINC01089*, *MEG3*, *PVT1*, *TINCR* and *UCA1*. According to Lnc2Cancer, the 98 lncRNAs are functional in mechanisms governing cell survival (n=48), cell growth (n=44), metastasis (n=37), EMT (n=25), drug resistance (n=12) and recurrence (n=8) (Table S9). From the 40 circulating lncRNAs in Lnc2Cancer (Table S9), detected in the plasma of breast cancer patients, we recovered 16, including the well-known biomarkers *GAS5* and *HOTAIR* [49].

Diermeier and collaborators [26] identified murine lncRNAs overexpressed in breast cancer samples (named mammary tumor associated RNAs, MaTARs), prioritizing for future studies a subset of 30 MaTARs conserved in humans. We identified four of these MaTARs in the normal breast: *ARPC1B-PDAP1*, *EMSLR* (*RP11-132A1.4*), *HOTAIRM1* and *TTC39A-AS1* (Table S9). As an example of the relevance of these genes for future research, ASO-mediated knockdown of *TTC39A-AS1* (MaTAR18) was shown to lead to impaired branch development in mammary organoids [26].

### C) *Annotated Seurat-assigned markers of L-clusters*

Fifty-six Seurat-assigned NB-lncRNA markers were previously annotated in the genome or noncoding gene databases. A few examples are general basal marker *NCF4-AS1* (DN1070C0G3I1; marker of clusters L1, L6 and L7), stem/progenitor marker *LINC02019* (DN24378C0G1I6; marker of clusters L3 and L5) and luminal progenitor marker *PXN-AS1* (DN90647C1G1I2; marker of clusters L0 and L2). Also, *ELDR* is an annotated lincRNA near basal marker *EGFR* [50] with an intronic basal marker NB-lncRNA (DN121756C0G1I2; marker of clusters L6 and L7). Annotated lncRNA markers were also found for A-clusters. Almost 8% of all Seurat-assigned markers (n=258) were not protein-coding (Table S13). Of the 23 lincRNAs markers, two (*LINC01060* and *LINC00342*) are general markers of luminal and basal cell types. *LINC00993* is a specific marker of the luminal mature cluster (A2) and a breast-specific lncRNA [51] located near *ANKRD30A*, a well-known luminal marker.

Several breast cancer-related lncRNAs were identified as Seurat-assigned markers of normal breast cell subpopulations. For example, *CARMN* (DN405C0G2I1), a general basal marker of clusters L1, L6 and L7, promotes cisplatin sensitivity in basal breast cancer [52]. *LINC01094* (DN19901C0G1I3), *MEG3* (DN4816C0G1I10) and *HOTAIRM1* (DN126378C0G1I5) are also Seurat-assigned basal markers, of clusters L6 and L7. *SNHG29* (DN1842C0G2I8) is a general luminal marker of clusters L0, L2 and L4 known to regulate senescence in breast cancer cells [53,54]. In agreement with observations made on A-cluster markers, two *LINC00993* isoforms are Seurat-assigned luminal mature markers of L-clusters (DN4775C0G2I12 and DN4775C0G2I13; cluster L4). Finally, luminal marker *LINC01151* (DN20235C2G3I1; clusters L0 and L4) has been implicated in breast cancer metastasis [55].

### SUPPLEMENTARY: SINGLE-CELL RNA SEQUENCING ANALYSIS

#### A) *Publicly available single-cell RNA sequencing data of the normal breast epithelium*

We used scRNAseq data from the normal breast cell atlas produced by Nguyen *et al* [56]. Briefly, authors sorted cells in populations by FACS flow cytometry using antibodies for CD31, CD45, EpCAM and CD49f, processed sorted samples either in the microfluidics-enabled platform (Fluidigm C1) or droplet-enabled platform (10x Genomics) and sequenced cells in Illumina HiSeq equipments (see original publication for details). Due to differences in the technologies, while the Fluidigm platform generated ~1.6 million reads per cell, the 10x Genomics platform generated an average of 50,000 reads per cell. Data from both platforms were downloaded from the SRA (PRJNA450409), including 867 cell files for individuals 1 to 3 (Fluidigm C1) and 4 samples, for individuals 4 to 7 (10X Genomics), which together comprise 24,646 cells. We decided to analyse the microfluidics-enabled scRNAseq data, reasoning that the greater sequencing depth would allow us to assess a more comprehensive set of lncRNAs.

#### B) *Methods for single-cell clustering with Seurat*

##### 1. Transcript sets used for single-cell clustering

We compared the cell clusters obtained based solely on expression of the NB-lncRNAs (L-clusters) with those obtained using only genes in the GENCODE-annotated transcriptome (A-clusters). To address specific questions, we also investigated expression of GENCODE-annotated confirmed lncRNAs (C-clusters) or protein-coding genes (P-clusters) and the set of annotated genes which are not protein-coding or confirmed lncRNAs (O-clusters). Finally, when the expression of lncRNAs and protein-coding genes needed to be assessed simultaneously, we used a comprehensive set of all GENCODE-annotated transcripts merged with the NB-lncRNAs (M-clusters).

##### 2. Initial assessment of cell types

We designed a script-based pipeline to process the data using a set of tools, which we made available through GitHub. Briefly, we used RSEM v.1.3.1 [57] to generate read counts based on Bowtie2 v.2.2.9 [11] alignments and the annotation GFF files of each transcript set described in '1'. Count matrices were imported to Seurat v.4.0.4 [58] using Melange v.0.1.0 (website reference[1]). For the GENCODE-annotated set of transcripts, cells with less than 900 detected genes were excluded by the quality control filtering, as well as all genes not detected in at least 3 cells after filtering. For the remaining transcript sets (which lack protein-coding isoforms), the minimum limit was lowered to 300 genes per cell, with every other parameter kept the same. Preliminary heatmaps based on the first principal components were plotted using Seurat's *DimHeatmap*.

##### 3. Dimensionality and resolution

After assessing principal components, we analysed the data using 'straw' and 'elbow' plots to define the dimensionality to be used for further clustering. *ElbowPlot* ranks principal components using the fraction of variance they each explain and *JackStrawPlot* compares the distribution of feature p-values for each principal component with that of a uniform distribution, characterizing the most significant ones as those enriched in low p-value features. Both graphs were manually inspected to define the most appropriate number of dimensions for clustering cells by UMAP based on each individual transcript set. The resolution is a parameter controlling the granularity of cell clustering in scRNAseq analyses. Increasing the resolution can lead to more informative clustering, but if set too high it can result in biologically meaningless results. We used a novel method named ClusTree v.0.4.4 [59] to thoroughly assess the effect of different resolutions (ranging from 0.2 to 1.2) and decide which to use in our analyses. Clustering trees (results generated by ClusTree, Figure S6) are by themselves a source of relevant information, especially on cell hierarchy, since they depict how 'superclusters' are subdivided into clusters and which 'subclusters' would result from an increase in resolution. The table below presents the values set for dimensionality and resolution in each clustering experiment. After careful examination we set the resolution at 0.5 for the analyses of A-clusters and M-clusters and increased the resolution to 1.0 for L-clusters, to enforce the separation of clusters L3 and L5, which clearly represented different cell populations.

| Experiment | Transcript set | Cluster name | Dim | Resolution | Min. features/cell |
|---|---|---|---|---|---|
| 1 | GENCODE-annotated, all genes | A-clusters | 9 | 0.5 | 900 |
| 2 | GENCODE-annotated, confirmed lncRNAs only | C-clusters | 6 | 0.5 | 300 |
| 3 | GENCODE-annotated, excluding protein-coding and lncRNA genes | O-clusters | 8 | 0.5 | 300 |
| 4 | GENCODE-annotated, protein-coding only | P-clusters | 8 | 0.5 | 900 |
| 5 | NB-lncRNAs | L-clusters | 6 | 0.8 | 300 |
| 6 | NB-lncRNAs merged with all GENCODE-annotated genes | M-clusters | 10 | 0.5 | 900 |

**4. Cell clustering**

After filtering performed as described above, we normalized counts per cell, performed logarithmic transformation and identified the 2,000 most variable genes. Seurat *LogNormalize* function divides feature counts for each cell by the total counts for that cell and multiply by the scale factor (which we set to 10,000), then performs natural-log transformation using log1p. *FindVariableFeatures* is used to identify a user-defined number of features (which we set at 2,000) characterized as outliers in relation to the mean. We then performed a linear dimensional reduction before plotting preliminary heatmaps to visualise the behaviour of principal components.

**C) *Results of single-cell clustering with Seurat***

**1. Major coding and noncoding markers of basal and luminal cell types**

Preliminary heatmaps were based on 200 'most extremely different' cells and the top 30 genes that most define these (in balance, i.e. 15 genes characteristic of each cell pool) sorted by principal component scores. For A-clusters, a clear separation between luminal and basal cell populations is obtained by these top genes, as expected, with well-known luminal markers (*KRT15, LTF, PI3, SLPI, WFDC2*) and basal/myoepithelial markers (*ACTA2, CCL2, IL1B, KRT14, TAGLN*) making up one third of the total.

Analysis of M-clusters heatmaps revealed novel NB-lncRNAs within the top 30 most discerning genes (Fig. A). The two major luminal marker NB-lncRNAs are 3'UTR TLARs for PIGR

(DN8937C2G1I2) and *SOX4* (DN11567C1G1I3). *PIGR* is responsible for the traffic of immunoglobulin across epithelial cells and has been shown to be a specific marker of luminal breast cancer [60]. Another portion of the *PIGR* 3'UTR sequence was implicated in colon cancer in a study by Traicoff and collaborators [61], who postulate it would have a role in *PIGR* mRNA stability. Notably, the original samples in which the region was observed were of mammary glands from healthy donors. *SOX4* is a well-known marker for luminal (likely luminal intermediate progenitors) cell populations [62]. Among the major basal markers, there is a NB-lncRNA intronic to *SAMD5*, which has been shown to be present at basal cells of resting mammary tissue (website reference[2]) and to be absent from luminal regions of normal bile ducts [63]. Another basal marker NB-lncRNA is a TALR from the 3'UTR of *CCND2*, which is a well-known basal marker [62].

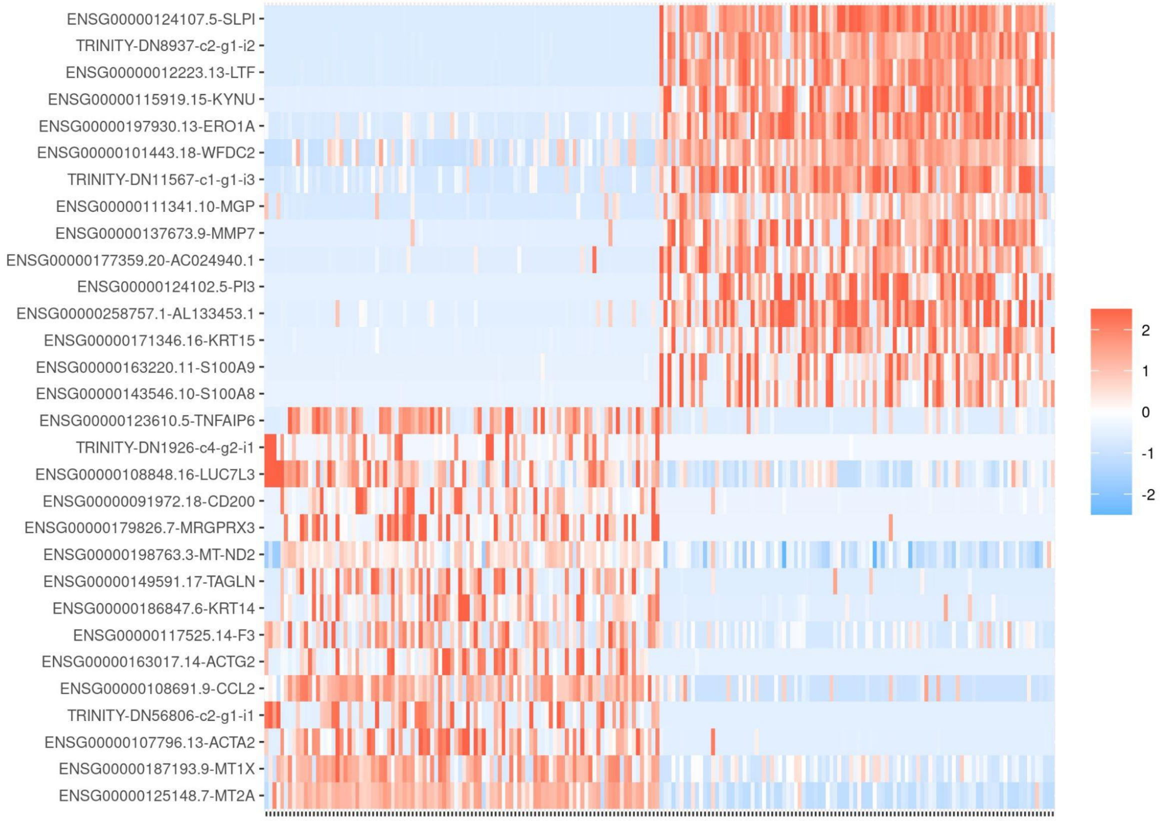


**Fig. A:** Seurat's heatmap of the top 30 most discerning genes between the two main cell groups, showing four NB-lncRNAs among the Seurat-assigned markers. The bar on the right represents expression levels.

### 2. Top 100 most variable genes across cells

We retrieved the top 100 genes with highest cell-to-cell variation observed in M-clusters. Sixteen of these were NB-lncRNAs: antisense intronic to *KMT2E* and *PSG9*; 3'UTR TALRs for *CCND2*, *FGFR1*, *TOR1AIP1*; sense intronic to *COG5*, *DRAM1*, *EPAS1*, *GLS*, *PSG5*, *PSMA3*, *PPP1R12B*, *RNMT*, *RP11-125D12.3*, *SH3PXD2A*, *TTC17*. Amongst the protein-coding counterpart, we found basal (*ACTA2, KRT14, KRT15*), luminal (*CYP24A1, SAA2, SLPI*) and other marker genes (epithelial marker *KRT17*, milk protein *PI3* and endothelial marker *SELE*).

### 3. Technical notes on data from the original publication

Nguyen *et al*. [56] reported only the three generic clusters (basal, luminal progenitor and luminal mature), based on data acquired in the Fluidigm C1 platform. Even at lower resolution (0.2) our method generated six clusters (two basal, two luminal mature, one luminal progenitor and one undefined cluster). Although we mostly recapitulated the methodology used in the original publication, some differences may account for the higher number of clusters, for example using a different dimensionality, using GENCODE annotation (instead of ENSEMBL) and clustering by UMAP (instead of t-SNE).
After cell sorting, authors have added labels ("BAS" or "LUM") to the files deposited to SRA, providing a way to confirm the broad identity we assigned to clusters. In total, there are 369 luminal and 498 basal cells in the samples (Table S12).

**4. Markers from laboratory applications**
Recently, Gusterson and Eaves [64] reviewed markers used in laboratory experiments to characterise and/or isolate different human mammary cell types. Their list is composed of genes that can be used in immunohistochemistry and other techniques. We assessed the distribution of these markers across cells in A-clusters and concluded that these alone are not sufficiently powerful to provide a clear separation of cell types and additional markers are needed to characterize cell types in the mammary epithelium.
Cytokeratins used to characterise breast cells by immunohistochemistry include myoepithelial (*KRT5, KRT6, KRT14*) and luminal (*KRT8* and *KRT18*) markers. Myoepithelial keratins *KRT5* and *KRT14* were found as markers of clusters A0 (basal) and A6 (stem-like), while *KRT6B* is a marker of clusters A0 and A1 (luminal). Both *KRT8* and *KRT18* were found as markers of clusters A1 and A2 (luminal), as expected, but not A4. *ITGA6*, used to prominently stain both basal and luminal cells, was found as a marker of cluster A3 (stem-like) only. *MME* and *CD44* are both present at higher levels on the surface of basal/myoepithelial cells and, while the former was found as a marker of cluster A6, the latter was found in cluster A3. Expression of basal marker *ACTA2* is significantly higher in clusters A0 and A6, with the former also exhibiting the highest *EGFR* levels. For luminal populations, *EPCAM* and *MUC1* are prominent staining agents for luminal cells in general and also *KIT*+ luminal progenitors. While *EPCAM* was confirmed as a marker for clusters A1 and A2, *MUC1* was a marker in all luminal clusters, A1, A2 and A4. KIT was only found as a marker of cluster A4. *PROM1* is higher on the surface of luminal cells and was observed on clusters A1 and A4.

**5. Clusters of heterogeneous cell subpopulations**
According to analysis of cluster trees (Figure S6), clusters A0 and A1 could be further subdivided into two populations each (by increasing the clustering resolution), indicating their higher cell heterogeneity (noteworthy, these are the most populated clusters). According to analysis performed comparing Seurat markers to markers from Pal *et al* [65], cluster A1 is a luminal progenitor but with significant overlap with luminal mature markers (p-value = 6.0e-10), indicating a possible intermediate nature. Cluster A0 is a basal 'supercluster' which we showed was subdivided into clusters L1, L6 and L7 based on NB-lncRNA expression. Increasing the resolution while generating A-clusters led to a segregation of this 'supercluster' in two clusters (Fig. B), one has >80% of its cells in L1 and the other has >80% of its cells in L6+L7. The differences between L1 and L6+L7 are reflected in the obtained trajectory, which placed L1 as a separate branch, originating directly from L3, while L6+L7 are part of another branch, which passes L5 after originating in L3 (Fig. 7d).

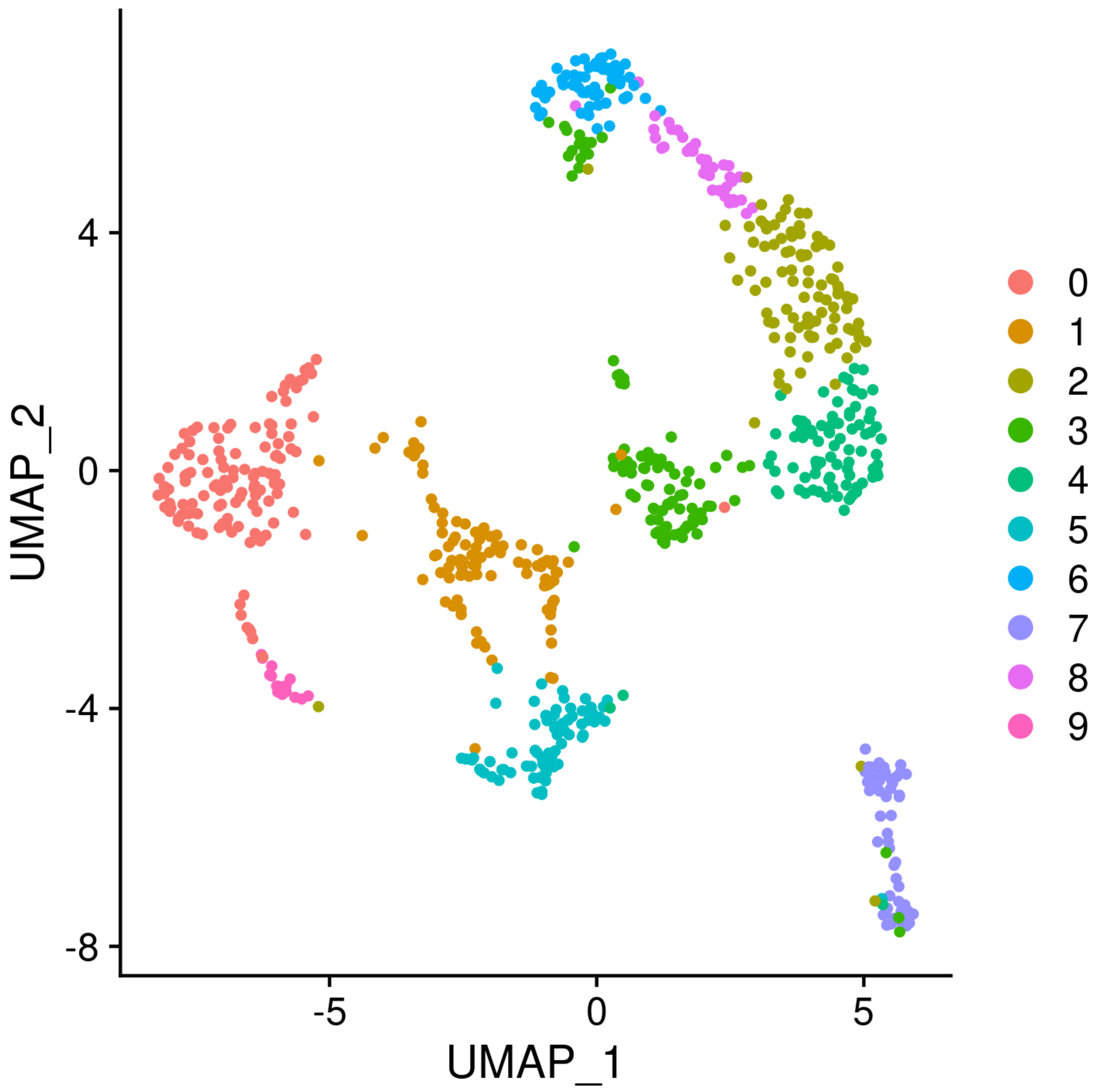


**Fig. B:** UMAP of A-clusters obtained after increasing the clustering resolution in Seurat. The experiment led to the subdivision of the heterogeneous basal cluster, but in only two subclusters (2 and 3 above).

**6. Clusters A5 and A7 likely include dead cells**

Since marker genes were not sufficient to characterize clusters A5 and A7, we investigated the 300 genes with highest average expression in each cluster (by calculating the average expression level of each gene across all cells and selecting the top 300). First, we noticed the same 14 (out of 37) human mitochondrially encoded genes within the top 20 most highly expressed of each cluster. In a scRNAseq experiment, this can indicate cell death. For comparison, all other clusters have only one or two such genes in their entire marker set. Although filters were in place to try and eliminate these cells from our analyses, a percentage of dead cells may still be present. Nevertheless, these genes make up only ~20% of all markers for each of the two clusters. The 300 most highly expressed genes of each cluster confirm their basal subtype (p-value 0.01 and 2.0e-11 for cluster A5 and A7, respectively compared with basal marker lists from Pal *et al* [65]). This is corroborated by the fact that these clusters are spatially close to cluster A0 and ClusTree shows cluster A7 derives from the same parental cluster as A0. Basal markers that are highly expressed in both clusters A5 and A7 include

*CALD1, CAV1, COL17A1, F3* and *ITGB1*. In addition, cluster A7 has high expression of TAGLN and SERPINB5 and cluster A5 of *A2M, ITGA2, KRT17* and *MYLK*. Interestingly, the main difference between A-clusters and M-cluster was that a subset of 20 cells from cluster A3 were placed with cluster A5, which was consequently renumbered as cluster M3 given the higher cell number. This could be a result of dead cells being properly separated from their live counterpart, although no clear difference in total number of mitochondrial-derived reads was observed in these 20 cells. Notably, while cells in cluster A5 are mainly absent from L-clusters (for failing filters), cells of cluster A7 cells remain and are scattered between basal clusters L5 and L6).

**7. Cluster characterisation based on protein-coding gene targets of NB-lncRNAs**

We listed the FEELnc-assigned protein-coding gene partners of marker NB-lncRNAs for each L-cluster and compared them with the list of 359 literature markers. Cluster L0 has NB-lncRNA markers associated with luminal marker genes such as *LTF*, *DBI* and *S100A6* (~65%). Cluster L1 NB-lncRNA marker targets include basal genes *ITGA6* and *TIMP3* (~20%) as well as immune, stem-cell and matrix marker genes (each also contributing with ~20% of the total). Cluster L2 has NB-lncRNA markers near luminal marker genes *ANKRD30A*, *DBI*, *IRX5* and *LTF* (~60%). Protein-coding partners of cluster L3 NB-lncRNAs markers are mainly stem-cell or prepubertal cell markers (~66% including *CD34*, *JAG1*, *TXN*, *CAMK2D* and *NCAM1*). Cluster L4 is more heterogeneous, with NB-lncRNA markers nearby luminal (*ANKRD30A* and *S100A6*) and basal (*COL4A1*, *IGFBP3* and *SOX9*) markers. Cluster L5 has no known protein-coding markers nearby its NB-lncRNA markers. Cluster L6 has NB-lncRNA markers near known basal (*EGFR*, *ITGA6*), general epithelial (*ALCAM*, and *CDH1*) and luminal (*ELF3* and *LTF*) markers and cluster L7 near immune (*CCL2* and *ANPEP*), stem-cell (*IGFBP4*) and matrix (*FGF2*) marker genes.

**8. Exemplary NB-lncRNA markers of interest**

In the basal subpopulations, a novel NB-lncRNA intronic to *SNRK* is a marker of all basal clusters (DN16110C0G1I1; clusters L1, L6 and L7). *SNRK* promotes breast angiosarcomas [66]. Another novel NB-lncRNA (DN390900C0G1I1) is a marker of basal clusters L1 and L6 intronic to *ITGA6* (CD49f), a gene commonly associated with basal populations. A lincRNA near *SATB2* is a marker of basal cluster L6 (DN21900C0G1I1; Fig. 4). *SATB2* is an interacting partner of *BCL11B*, which governs quiescence by preventing the exhaustion of stem-cells through expression of the cell cycle inhibitor *CDKN2A* [67]. Two NB-lncRNAs (DN33019C0G1I7 and DN33019C0G1I15) are markers of cluster L7 localized near *EDN1*, which is a well-known endothelial marker. A 3'UTR TALR (DN22692C2G1I1) of *CAMK2D* (a prepubertal marker gene) is a marker of clusters L3 (stem-like), L1 and L7 (basal). Two annotated NB-lncRNAs (DN5838C0G2I13 and DN87246C0G1I6), specific markers of the basal cluster L7, are respectively antisense to *TTC28* and *FOXG1*. A FOXG1-associated lincRNA (LINC01551) is a marker of the correspondent A-cluster (basal cluster A0).

A novel (unannotated) NB-lncRNA is a luminal progenitor marker (DN3664C0G3I1; clusters L0 and L2) located near *LTF* which is a known luminal progenitor marker. Another novel NB-lncRNA, intronic to breast stem-cell marker *TXN* is a stem-like cluster marker (DN19566C1G2I1; cluster L3). A novel NB-lncRNA intronic to *CCDC25* is a marker of all three luminal clusters (DN225116C0G1I1; clusters L0, L2 and L4). *CCDC25* is a breast cancer metastasis promoter [68,69].

**9. Ubiquitously expressed versus compartmentalised transcripts**

We divided transcripts into those expressed in more (ubiquitous expression) or less (restricted expression) than ~1/3 of all cells (241/741 cells or 32.5%). To directly compare transcript counts of protein-coding genes and NB-lncRNAs, we assessed their expression in M-clusters, in which both are quantified simultaneously. In this clustering experiment, a total of 17,667 protein-coding genes, 13,900 NB-lncRNAs and 10,327 annotated and confirmed lncRNAs were detected in at least one validated cell. There were 4,586 ubiquitously expressed protein-coding genes (25.9%), 602 ubiquitously expressed confirmed lncRNAs (5.8%) and 121 ubiquitously expressed NB-lncRNAs (0.9%). We then separated genes of restricted expression into those expressed in 12-40 cells and 41-241 cells and calculated their median TPM counts in each interval. For protein-coding genes, the expression range is between 11 and 13 TPM, while for NB-lncRNAs is between 19 and 23. Median expression for annotated and confirmed lncRNAs ranged between 7 and 14 TPM.

Nearly half (2,168) of the 4,586 ubiquitously expressed protein-coding genes have been previously shown to be of housekeeping function by the HK and/or HRT databases [70] and [71] which together contain ~4,500 housekeeping genes; Table S8). This provides proof-of-concept that widespread protein-coding gene expression is significantly (p-value of overlap ~0) correlated with housekeeping functions. Proportionally, the number of NB-lncRNAs likely involved with housekeeping functions is nearly the same as protein-coding genes, with 56 of the 121 ubiquitous lncRNAs having at least one housekeeping partner (p-value 5.4e-31). Interestingly, for 26 of the 56 lncRNAs, at least one co-expressed housekeeping gene is in the same chromosome.

Examples include NB-lncRNAs near *MKLN1*, *PSMD7* and *SYNCRIP* (DN12454C2G3I3, DN42909C1G1I2 and DN98365C2G1I1), and those antisense to *IAH1* and *TIMM44* (DN82157C1G1I1 and DN49627C1G1I1). See Table S15 for a comprehensive list.

**10. NB-lncRNAs define subcluster regions**

Since we showed NB-lncRNA expression to be more compartmentalised than that of protein-coding genes, we reasoned that they were not homogeneously distributed across each cluster, but rather segregated to subsets of cells. Such expression patterns should result in increased specificity indices, if clusters were further subdivided by increasing the resolution parameter of Seurat's *FindClusters* function. We therefore re-clustered cells, gradually increasing the resolution from 0.2 to 1.0 and calculated specificity indices for protein-coding transcripts and NB-lncRNAs. For protein-coding transcripts, the overall increase in specificity was of only ~10% (from 3.6 to 4.0) and not all resolution increments led to higher average specificity indices. For NB-lncRNAs, however, the overall increase was of ~60% (from 3.4 to 5.4) and every increment in resolution led to higher average specificity indices. Because the sum of specificity indices in all clusters for a given transcripts is always 1.0, increasing the number of resulting clusters by increasing the resolution would have a general effect of reducing the index in every cluster. To account for this, we normalised the specificity indices by the inverse of the number of resulting clusters. We also investigated the number of transcripts for which specificity indices were either the same or higher as resolution was increased. From 17,778 protein-coding genes, 2,330 (13.1%) presented this behavior, while from 14,595 NB-lncRNAs, 4,327 (29.6%) did.

#### ***Website References:***

1: https://rdrr.io/github/rahuldhodapkar/melange

2: https://api.research-repository.uwa.edu.au/ws/portalfiles/portal/12342713

**SUPPLEMENTARY: REPLACEMENT OF SCENT GENE NETWORK FOR POTENCY ASSIGNMENT**

**A)** ***Retrieve lncPathway data***

LncRNAs2Pathways [72] is a method to assess the contributions of lncRNAs to biological pathways. The method relies on the placement of the lncRNAs in a gene interaction network, based on expression correlations between lncRNAs and coding genes. The algorithm is freely available as an R-based tool (website reference[1]). The underlying coding-noncoding gene network (CNC network) was constructed by integrating RNAseq data and protein-protein interaction data. Authors collected 28 human RNAseq datasets covering a wide range of experimental and physiological conditions from NCBI and SRA (Supplementary Table S1 in [72]). Annotations of all human lncRNAs and protein-coding genes were downloaded from GENCODEv22. Based on expression data generated from the RNAseq experiments, authors identified 114,006 co-expression relationships of coding-coding, coding-lncRNA and lncRNA-lncRNA genes. Authors then integrated gene co-expression data with protein-protein interactions retrieved from the Human Protein Reference Database, the Database of Interacting Proteins, the Molecular INTeraction database and Reactome. Gene co-expression relationships were merged with protein-protein interactions to construct a CNC network with 28,613 nodes and 295,698 edges.

We recovered the annotation of 28,202 such genes, 40% of which were not protein-coding. For comparison, in both internal SCENT [73] networks (net13Jun12.annotation and net17Jan16.annotation), less than 50 genes are not protein-coding. Below are the gene classes that form the CNC network of LncRNAs2Pathways.

| Gene Class* | Number of Genes |
|---|---|
| protein_coding | 17582 |
| lincRNA | 4518 |
| antisense | 4125 |
| TEC | 713 |
| sense_intronic | 635 |
| processed_transcript | 393 |
| sense_overlapping | 151 |

* showing only classes with >100 elements

A summary of the code used to generate the final table used as input for SCENT is contained in Script File 7. Genes in the CNC network were identified by a mixture of gene names (17,599 entries) or Ensembl IDs (10,992 entries). For consistency, we mapped gene names to GENCODEv36 gene IDs. Unfortunately, some gene names have been revised and changed recently. To account for these updates, we used HUGO gene annotations. From the 17,599 gene names, 16,537 were directly identified in GENCODEv36 and another 1,040 were retrieved after their updated name was retrieved from HUGO (bringing the total to 17,577, with 22 missing). From the 10,992 Ensembl IDs, 10,530 were directly identified in GENCODEv36 (with 462 missing). Most of the nearly 500 missing genes were further retrieved from Ensembl, 33 of which were also in GENCODEv36, bringing the total GENCODE-annotated genes to 28,140 (>98%). The list of gene names with GENCODEv36 IDs was supplemented with empty IDs, to account for all annotated genes.

**B)** ***Internal SCENT networks***

We used *bioDBnet* (website references[2,3]) to retrieve the Entrez IDs of GENCODEv36 genes. All 60617 genes were searched, with 25,492 being mapped and 35,083 failing to be mapped. There are 2 networks available in SCENT: *SCENT_net13Jun12.matrix*, has 8,434 genes and *SCENT_net17Jan16.matrix* 11,751. The great majority (>99%) of all genes are protein-coding.

#### *Website References:*

1: https://cran.r-project.org/web/packages/LncPath
2: https://biodbnet.abcc.ncifcrf.gov
3: https://biodbnet.abcc.ncifcrf.gov/db/db2dbRes.php

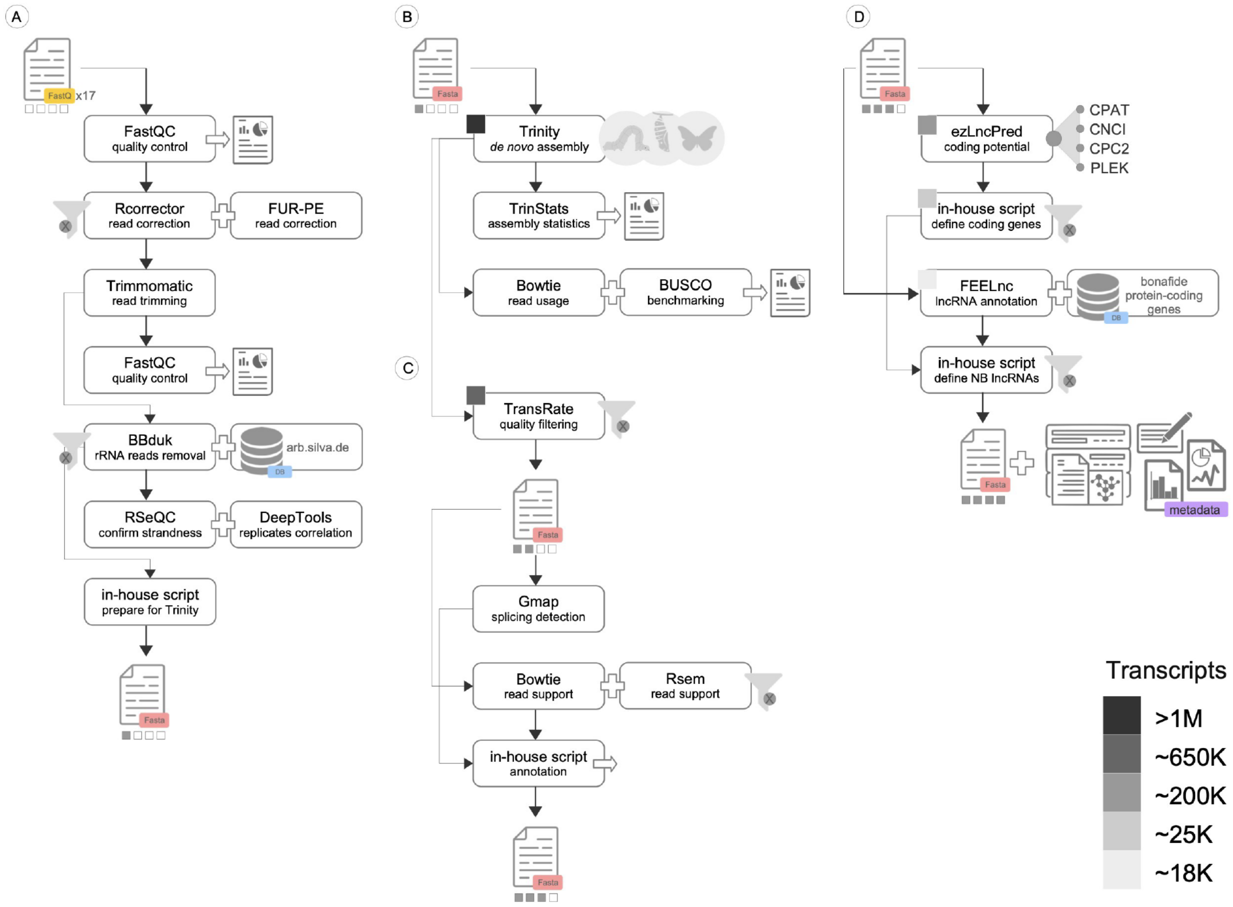


**Fig. S1 Computational pipeline for the identification of NB-lncRNAs.** Details on the multistep computational pipeline designed for *de novo* transcriptome assembly. Each main step (A-D) is expanded from Fig. 1a to show the intermediate steps and the tools used to perform each task. The approximate number of transcripts left after each filtering routine is shown as boxes in grayscale.

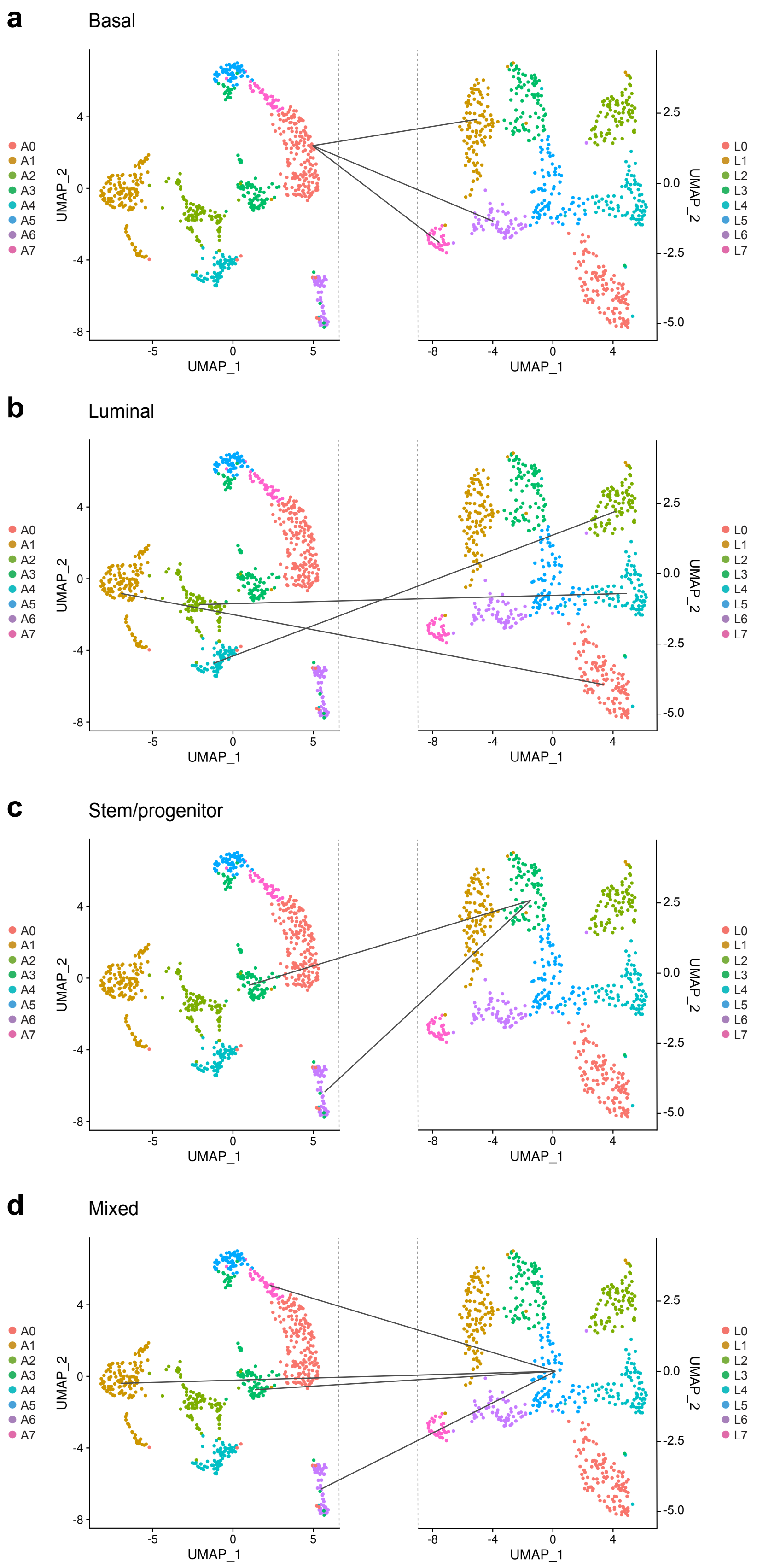


**Fig. S2 Correspondence between L-clusters and A-clusters, based on cell composition.** Cluster correspondence was defined based on the cell composition of each A-cluster (left) and L-cluster (right). Corresponding clusters are connected with black lines. **a** Cells in cluster A0 are spread in clusters L1, L6 and L7. **b** Cells in clusters A1, A2 and A4 are mostly (>80%) assigned to respectively clusters L0, L4 and L2. **c** Most cells in clusters A3 and A6 are combined into cluster L3. **d** Smaller proportions (15-25%) of cells from clusters A1, A3, A6 and A7 are combined into cluster L5.

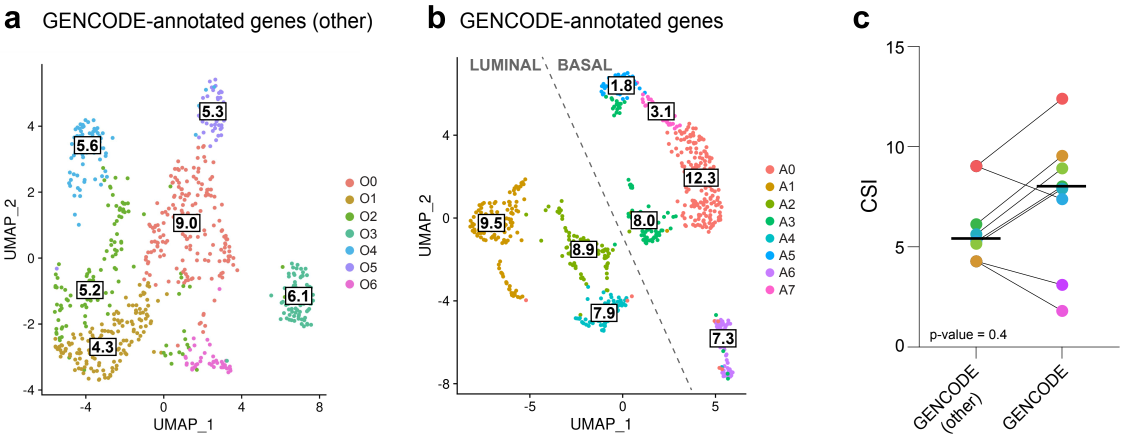


**Fig. S3 Higher cluster specificity is a characteristic of NB-lncRNAs. a** UMAP showing normal breast cell clusters obtained based on gene expression of annotated genes which are not protein-coding or confirmed lncRNAs (O-clusters), with their corresponding cluster specificity index (CSI). **b** UMAP showing normal breast cell clusters obtained based on GENCODE-annotated gene expression (A-clusters), with their corresponding CSI. **c** Dotplots showing the difference in CSI for corresponding clusters in **'a'** and **'b'**. Dots were colored according to the represented O-cluster or A-cluster, bold horizontal lines mark the average CSI for each gene set and the p-value (0.4; Fisher's exact test) shows the difference is not significant. As the CSIs of O-clusters were not normally distributed (according to the Shapiro-Wilk test), a Wilcoxon rank test was performed instead of a t-test.

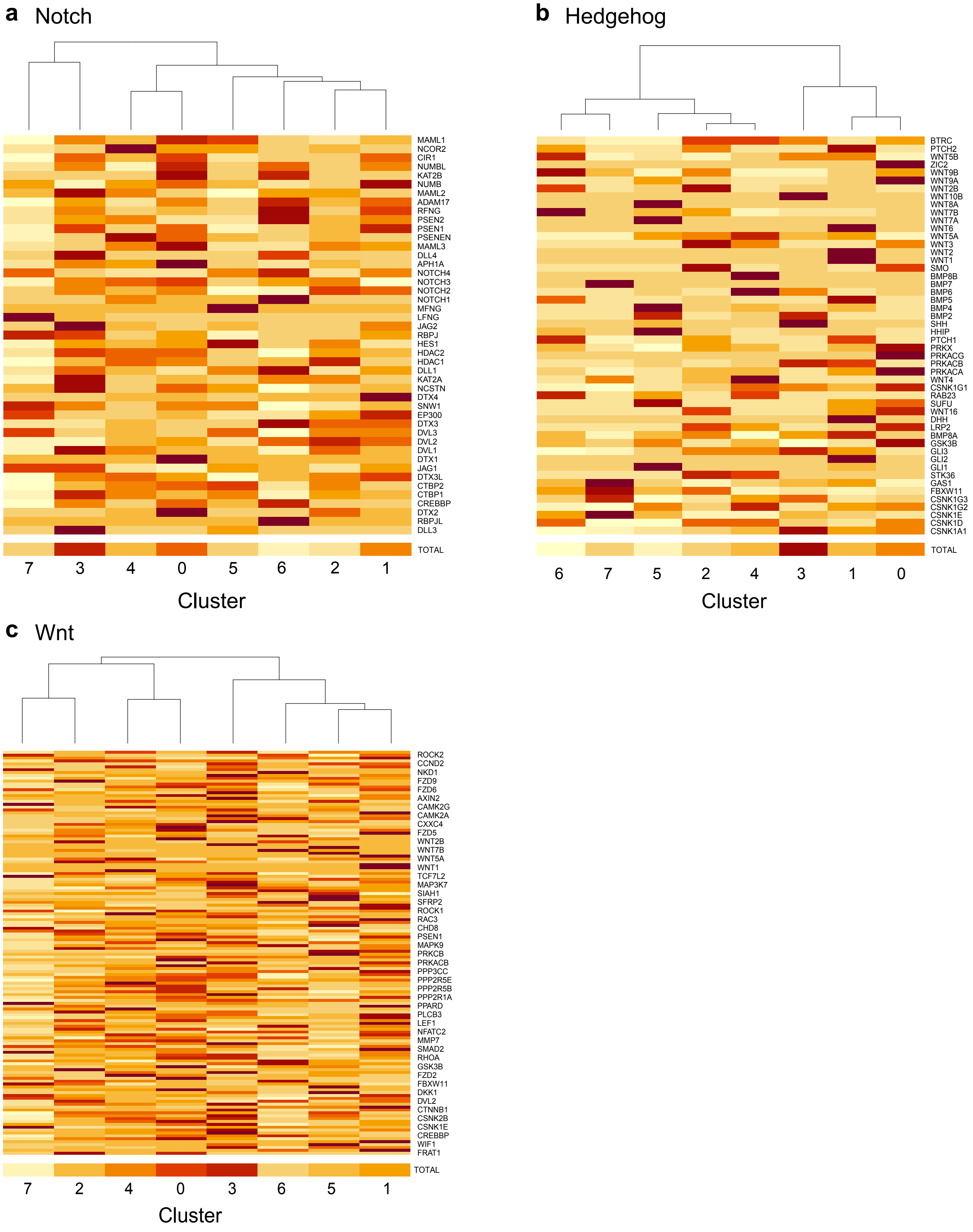


**Fig. S4 Notch, Hedgehog and Wnt signaling pathways are activated in cluster L3.** As a method to determine the L-cluster most likely harboring normal breast stem-cells, we assessed the expression of genes involved in the three main pathways associated with breast stemness, **a** Notch, **b** Hedgehog and **c** Wnt signaling. Based on the summarized gene expression, shown as the bottom row in each heatmap, cluster L3 was the one with highest overall expression of the genes of interest for each pathway.

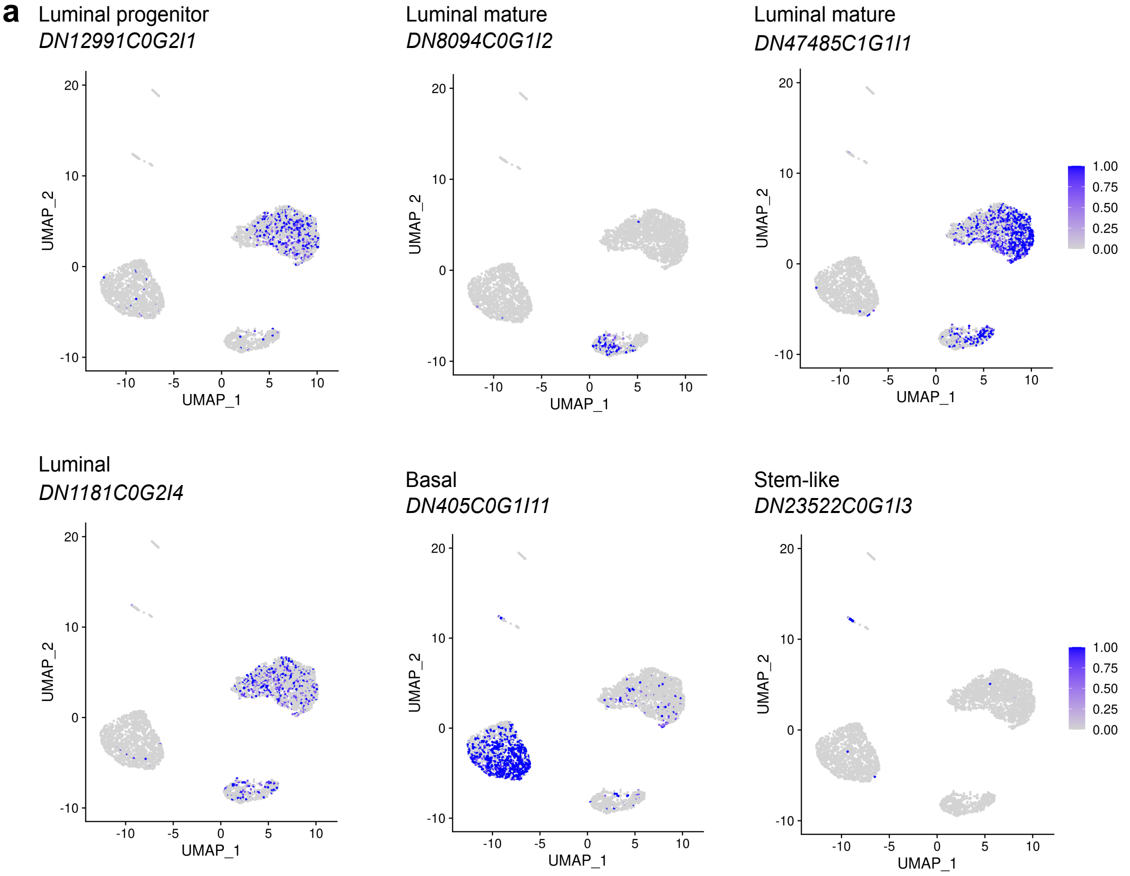


**b**

| GENCODE-annotated | | GENCODE-annotated + NB-lncRNAs | | |
|---|---|---|---|---|
| Cluster 7 | Cluster 9 | Cluster 9 | Cluster 10 | Cluster 11 |
| YBX1 | YBX1 | YBX1 | YBX1 | |
| ACAT1 | BTF3 | BTF3 | ACAT1 | FABP5 |
| FAU | CD44 | CD44 | FAU | GNG11 |
| RPL8 | GNG11 | IGFBP4 | RPL8 | HMGA1 |
| RPS18 | IGFBP4 | KLF4 | RPS3 | RPS3 |
| TCF4 | KLF4 | PROCR | RPS18 | TCF4 |
| | TXN | TXN | | TXN |
| | | ZEB2 | | |

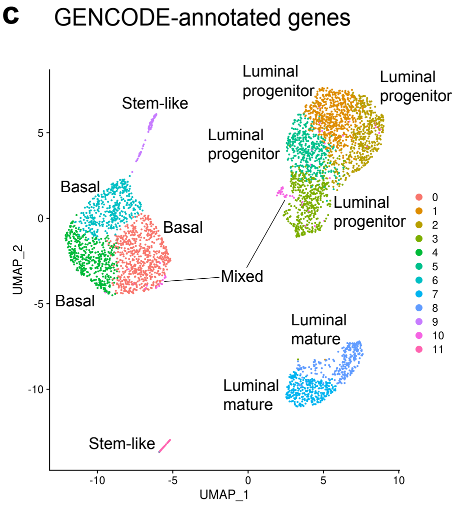


**Fig. S5 Seurat-assigned NB-lncRNA markers for cell clusters of 10x Genomics scRNAseq data. a** Expression patterns of identified NB-lncRNA markers for specific cell subpopulations in clusters computed for 10x Genomics scRNAseq data based on NB-lncRNA expression. **b** Known markers of stem cells in each of the clusters labeled as "putative stem-like". **c** Forced overclustering of 10x Genomics scRNAseq data based on expression of GENCODE-annotated genes also resulted in the subdivision of the luminal population into three clusters, but no additional stem-like clusters were observed.

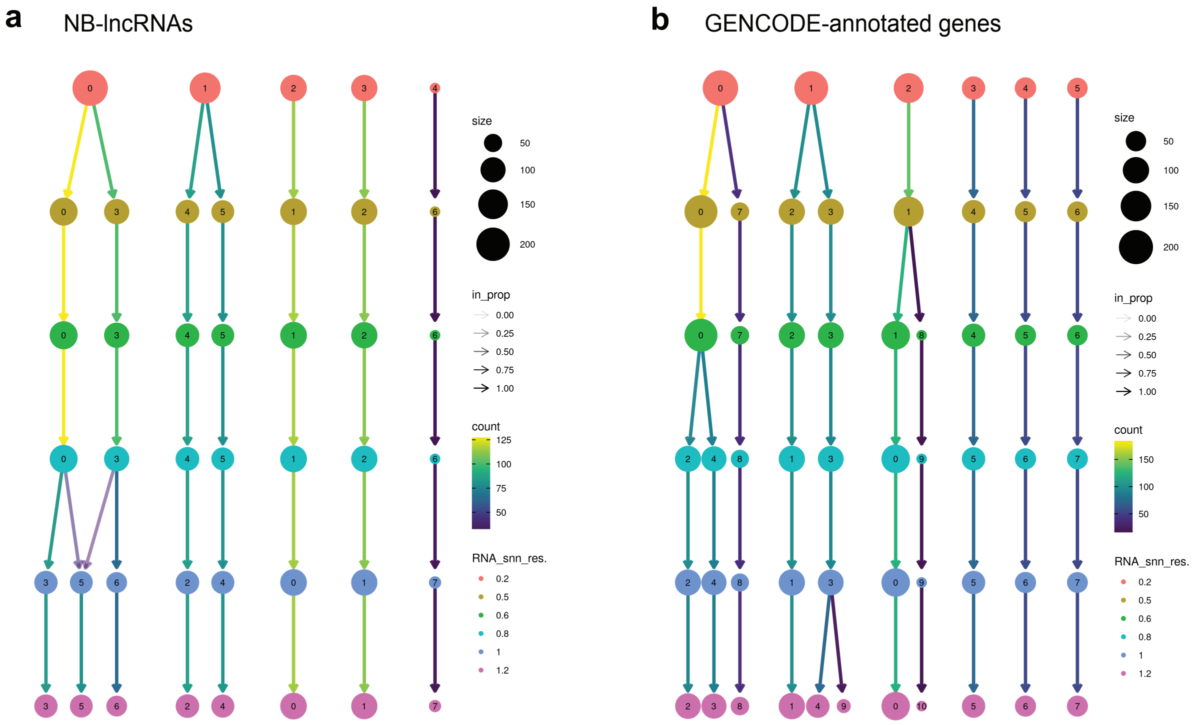


**Fig. S6 Clustering trees showing the relationships between clusters at various resolutions.** Clustering trees generated with ClusTree v.0.4.4[66] for clustering experiments based on the expression of **a** NB-lncRNAs or **b** GENCODE-annotated genes.
